# Spectral dependency of the human pupillary light reflex. Influences of pre-adaptation and chronotype

**DOI:** 10.1101/2021.05.28.446114

**Authors:** Johannes Zauner, Herbert Plischke, Hans Strasburger

**Author notes:** Corresponding author: (JZ). HP and HS are Joint Senior Supervisors.

## Abstract

Non-visual photoreceptors (ipRGCs) and rods both exert a strong influence on the human pupil, yet pupil models regularly use cone-derived sensitivity as their basis. This inconsistency is further exacerbated by the fact that circadian effects can modulate the wavelength sensitivity. We assessed the pupillary reaction to monochromatic light stimuli in the mesopic range. Pupil size for eighty-three healthy participants with normal color vision was measured in nine experimental protocols with varying series of continuous or discontinuous light stimuli under Ganzfeld conditions, presented after 90 seconds of dark adaptation. One hundred and fifty series of stimulation were conducted across three experiments, and were analyzed for wavelength-dependency on the pupillary constriction amplitude (PCA), conditional on experimental settings and individual traits. Traits were surveyed by questionnaire; color vision was tested by *Ishihara plates* or the *Lanthony D15* test. Data were analyzed with generalized additive mixed models (GAMM). The pupillary constriction amplitude response is consistent with L+M-cone derived sensitivity when the series of light stimuli is continuous, i.e., is not interrupted by periods of darkness, but not otherwise. The results also show that a mesopic illuminance weighing led to an overall best prediction of pupillary constriction compared to other types of illuminance measures. IpRGC influence on PCA is not readily apparent from the results. When we explored the interaction of chronotype and time of day on the wavelength dependency, differences consistent with ipRGC influence became apparent. The models indicate that subjects of differing chronotype show a heightened or lowered sensitivity to short wavelengths, depending on their time of preference. IpRGC influence is also seen in the post-illumination pupil reflex if the prior light-stimulus duration is one second. However, shorter wavelengths than expected become more important if the light-stimulus duration is fifteen or thirty seconds. The influence of sex on PCA was present, but showed no interaction with wavelength. Our results help to define the conditions, under which the different wavelength sensitivities in literature hold up for monochromatic light settings. The chronotype effect might signify a mechanism for strengthening the individual’s chronotype, but demands replication in a controlled study.

## Introduction

The human pupil is of interest to various research fields, such as vision research, neurobiology, ophthalmology, and psychology. Pupil constriction and size are controlled by parasympathetic pathways, and dilation through sympathetic pathways. Pupillary reaction thus acts as a window into the autonomous nervous system and the pupillary reaction coincides with, or even predicts, human behavior. Diagnostic methods were thus developed around various aspects of the pupil’s behavior and a vast body of research is dedicated to the pupillary reaction to light. One of the most fundamental aspects in that context is the dependency of the pupillary light reflex (PLR) on the spectral composition of light.

Up until now, the literature is somewhat divided on what the appropriate spectral weighing function should be. Pupil models derived from research using white light stimuli, i.e., polychromatic light spectra, generally employ stimulus luminance or retinal illuminance as predictors. Watson and Yellott (1) in 2012 reviewed eight pupil models published between 1926 and 1999. They created a ninth model by incorporating elements of the previous ones. What is important in our case is that all these models, including the newly created one, work with standard photometric dimensions, and spectral calculations are thus based on the V(λ) weighing function, regardless of stimulus characteristics. However, the V(λ) function may not be the best-suited for predicting pupil size in all cases.

The V(λ) function peaks at around 555 nm wavelength. Yet studies using monochromatic light stimuli generally report a maximum sensitivity for PLR at around 480 – 510 nm. Early publications to this effect are notably by Wagman and Gullberg (2) in 1942, by Alpern and Campbell (3) in 1962, and the often-cited paper by Bouma (4) in 1962. These authors attributed the PLR’s spectral dependency mainly to the characteristics of rod photoreceptors, but possibly also to those of short-wavelength cone receptors. Adrian (5) in 2003 tried to explain the apparent blue shift in sensitivity, reporting that, with appropriate adjustments, stimulus luminance was sufficient for explaining the PLR. According to the paper, a V_10_(λ) spectral weighing function which is based on the larger 10-degree field should be used instead of the usual 2-degree field (on which V(λ) is based) which, according to the author, is valid for the often-used Ganzfeld conditions. Furthermore, Adrian argues, mesopic conditions should be adequately addressed by using V_eq_(λ) weighting functions as intermediaries between the photopic V_10_(λ) and the scotopic V’(λ) functions. With these adjustments, pupil size depended linearly on (adjusted) luminance. While Adrian (5) very likely did not know about the spectral-temporal changes of the PLR as found in later studies, his argument about the appropriate luminance function remains valid, even when it was not widely adopted.

With the discovery of intrinsically photosensitive retinal ganglion cells (ipRGCs) at the beginning of the 21st century [6], and their potential to drive pupillary constriction [7], researchers began looking for their influence in PLR experiments. In 2007, Gamlin et al. (8) made the ipRGC influence on pupilloconstriction evident in the macaque with pharmacological blockade of rod and cone photoreceptors. They also showed ipRGC dependency for the post-illumination pupil reflex (PIPR) of humans. Zaidi et al. (9), also in 2007, confirmed the human pupilloconstriction’s dependency on ipRGC sensitivity in a blind person who lacked an outer retina. In 2009, Mure et al. [10] found ipRGC-dependent pupilloconstriction in the PIPR of sighted humans and, more importantly, during more prolonged light stimulation than used in previous studies (five minutes, compared to a few seconds in earlier studies). In 2010, McDougal and Gamlin (11) investigated the contributions of individual receptors to the PLR, for stimulus durations between 1 and 100 seconds. They found that rods play an essential role for most stimulus durations, while cones contribute only minimally after just 10 seconds of stimulus onset. At about 18 seconds after stimulus onset, the ipRGC sensitivity curve already provided the most prominent contribution. In 2012, Gooley et al. (12) used stimulus durations from 2 to 90 minutes, showing for the PLR, among others, the dominance of shorter wavelengths at around 490 nm, compared to the brighter settings around 555 nm, especially for longer stimulus durations.

Spitschan (13) showed the inadequacy of using V(λ)-derived luminance instead of a melanopic weighing function, estimating the possible error for the stimulus calculation to predict pupil size at about one order of magnitude for typical white-light sources. A recent paper by Zandi et al. (14) looked at the prediction accuracy of the models of Watson and Yellott (1). They found that the prediction error was greatest for chromatic spectra, especially for longer exposure times. The authors go as far as calling the luminance-based approach obsolete in light of all the known receptoral inputs. Spitschan (13) was more moderate in his evaluation, claiming that the actual relevance would depend on the application’s circumstances.

In summary, by addressing the effects of time since light onset, the ipRGC’s influence on PLR has become evident for monochromatic stimuli [15]. These findings sparked the development of diagnostic methods [16] for assessing the health of the nonvisual pathway [17] and circadian system [18], and spotting specific pathologies early on, such as glaucoma [19] and diabetic retinopathy [20]. However, what is still lacking is knowledge about why the PLR seems to differ in spectral sensitivity between monochromatic light and polychromatic (white) light. The applicability of prior findings based on monochromatic stimuli may be limited for polychromatic or even monochromatic settings. Furthermore, there is still little research into differences of spectral dependency beyond comparing particular wavelengths, mostly those of the blue and red peaks. Most of the studies that use a broad range of wavelengths are based on a comparatively small sample size. The largest of the studies reviewed above had twenty-four participants [12], while the median is only four participants [2-5, 8-12]. While this is enough for investigating the fundamentals of the PLR’s wavelength sensitivity, exploring distributional relationships will require more participants.

We initially set out to take a comparatively large sample of participants for looking at the conditional spectral dependency of pupilloconstriction. Since the ipRGC influence on PLR is mainly present for long-lasting monochromatic light stimuli, we believed that, basically, pupilloconstriction would approximately follow the ipRGC sensitivity curve for a series of continuously applied monochromatic light stimuli that change in wavelength over several minutes. Preliminary measurements supported this view and further indicated a dependence on sex, the latter dependency also having some support in the literature [21]. We were further interested in the influence of chronotype and time of day (cf. [18, 22]); findings from Zele et al. (23) suggested that there is a circadian variation in the wavelength-dependency, as demonstrated by pupil reactions to red and blue stimuli. In our main experiment with a balanced design, wavelengths around the V(λ) function’s peak led to the highest constriction. Based on these findings, we designed an exploratory, second experiment, to test whether our findings held up with a changed stimulus series. The findings were indeed replicated, but only in the case when light remained on continuously. When light was discontinuous, i.e., when short intervals of darkness were introduced between the light steps, shorter wavelengths became more influential, in line with published literature. The newly introduced periods of darkness in the second experiment additionally allowed spectral analysis of the post-illumination pupil reflex (PIPR). In a third experiment, we therefore investigated (1) the spectral dependency of the PLR for longer-lasting light steps with short, intermediate periods of darkness, as well as (2) the PIPR for short light steps with more prolonged intermediate periods of darkness.

We believe our findings will help bridge the gap between the canonical divide of luminance-based versus short-wavelength-based pupil models by showing that the PLR’s response to monochromatic light stimuli can go both ways, depending on pre-adaptation to light. We also show the influence of chronotype on pupilloconstriction, depending on the time of day. Statistical analyses employ generalized additive models (GAMs), which to our knowledge have not been used in pupillometry before. They appear particularly well suited for disentangling the complex coaction of influences.

## Materials and methods

### Participants

We recruited a total of 83 young, healthy participants across three experiments through bulletin boards and announcements at the Munich University of Applied Sciences (45 females and 38 males; age: median 26 yr., range 18 – 36; chronotype score: 50 ± 10.6 on the Morningness-Eveningness Questionnaire [24]). All procedures were approved by the Ethics Committee of the Munich University of Applied Sciences. Exclusion criteria were age (above 39), issues of psychiatric or neuronal health, addiction diseases, regular intake of stimulants or sedatives, acute jetlag or shiftwork (during the past three months), ocular diseases, or epilepsy. Participants were instructed to refrain from caffeine intake prior to the experimental sessions. We further checked participants for normal color vision (Lanthony D15 test [25] or Ishihara plates [26]). Additional participants were tested but their results not included in the analysis [27]; these participants either only took part in the preliminary measurements (see below), did not fit the inclusion criteria, or were tested after the experiment’s cut-off date.

### Apparatus

The light stimuli were presented in a Ganzfeld dome setup (Light Dome Model *XE 509, Monocrom,* Stockholm). The light source was a 100-Watt Xenon lamp. Full-spectrum light (6000 Kelvin correlated color temperature) was filtered through a stepper-controlled monochromator and led into the dome’s interior by a diffusor which was positioned above and behind participants’ eyes and thus not visible during the experiment. Light onset and offset were controlled by a shutter mechanism, located between the monochromator and the diffusor. Shutter response time, i.e., the time between open and closed states, was about 100 ms. Changes in peak wavelength occurred at a speed of about 35 nm/s. An *Arduino MEGA-2560* embedded computer with custom-written software controlled the monochromator and shutter.

The setup allows generating monochromatic light, with spectral peaks at between 400 nm and 700 nm, and a full width at half maximum of 12 nm ± 1 nm (*Fig 1*). The radiant output from the light source is dependent on wavelength (a limitation of the apparatus) and irradiance increases by about one order of magnitude from 404 nm to 450 nm; above that and up to 690 nm, irradiance is comparatively constant at a level of −1,62 ± 0.04 log_10_(W/m^2^). To accommodate for that dependency, in particular the lower irradiance levels for the wavelengths at and below 450 nm, we included the log_10_ of the irradiance as a covariate in all statistical analyses when it was not already included implicitly, i.e., as following from illuminance. Measurements of spectral irradiance were taken at the eye level under the light dome. Measurements further included a human field-of-view restriction suggested by CIE Standard *S 026/E:2018* [28]. Therefore, the irradiance measurements represent the corneal level, as do the various illuminance calculations based on those measurements (*Fig 1*). Unobstructed measurements, i.e., *regul*ar irradiance measurements, were about 24% higher.

**Fig 1.**
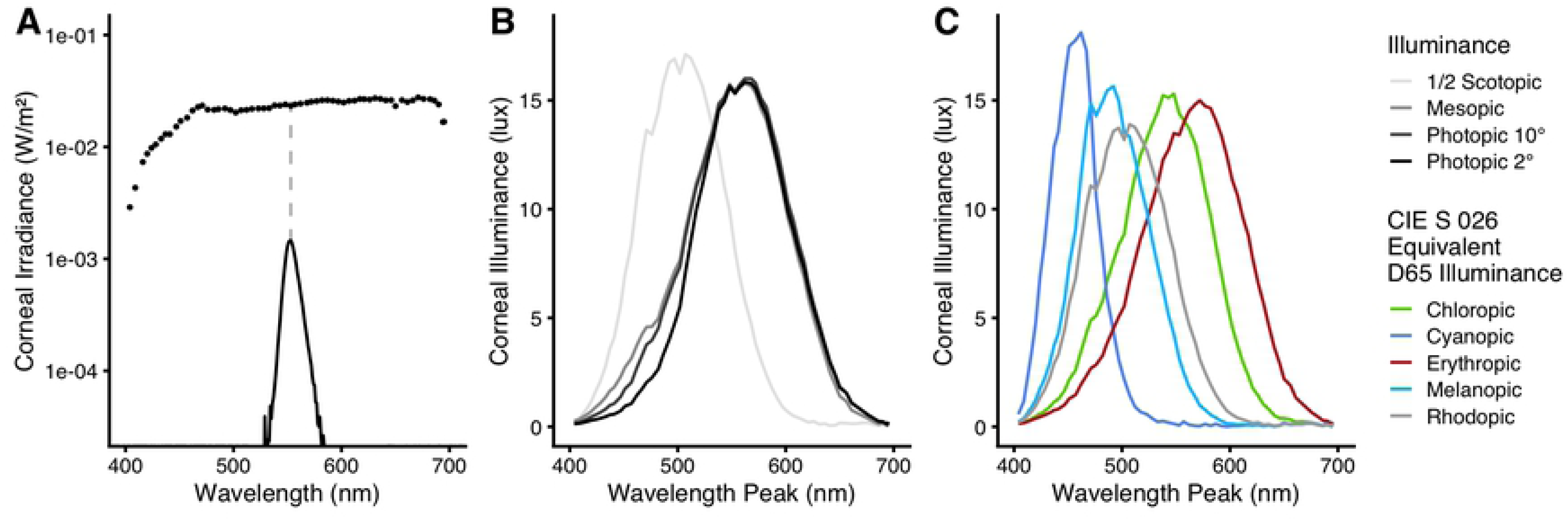
Spectral irradiance measurements and calculated illuminance values at the eye level. All displayed values are based on spectral irradiance measurements with a field-of-view restriction according to the CIE *S 026* standard [28]. In our case, these measurements are 24% lower than those of the unobstructed sensor diffusor. (A) Dots show the corneal irradiance (in W/m^2^) of exemplary steps at their respective peak wavelength. Each dot represents a monochromatic peak, similar to the spectral example distribution with a peak at 553 nm (the peak is indicated by the gray dashed line). (B) Corneal illuminance values for the monochromatic light steps at each peak wavelength. Besides the *standard* photopic CIE-1931 V(λ) weighing for the 2° observer (black line), illuminance values were calculated based on the weightings of V_10_(λ) (10° observer, photopic vision, dark grey), of V’(λ) (scotopic vision, light grey), and of V_eq_(λ) (mesopic vision, medium grey). To show the scotopic illuminance values along with the other curves, they are drawn at half their original value (light-gray curve). (C) Corneal alpha-opic equivalent-daylight illuminance for all receptor types according to the CIE *S 026* standard [28].

### Measurement equipment

An eye tracker (*Dikablis Professional Glasses*, *Ergoneers GmbH*, Munich) recorded the pupils by a dual infrared camera setup for simultaneous measurement of the two eyes, at a sampling rate of 60 Hz. Cameras in that setup are located at the end of flexible arms and are adjustable for an optimal view of the pupil and distance from eye level. Each camera has a resolution of 384 by 288 pixel. Camera pictures were analyzed in real time while recording through the *D-Lab 3.5* software (*Ergoneers GmbH*) on a connected personal computer. The software extracted several pupil parameters for each eye in pixel units, including pupil area, height, and width. These time-stamped variables were then exported to a comma-separated-value (CSV) file for later analysis. Spectral irradiance measurements with a 1-nm resolution were performed using a *JETI Specbos 1201* spectroradiometer (*JETI Technische Instrumente GmbH*, Jena; see *Fig 1*), with the *JETI LiVal V6.14.2* software running on a connected personal computer.

### Experimental Design

#### Preliminary measurements

In the relevant literature, the employed periods of dark adaptation (DAP) before light onset vary widely, if reported at all. They range from no-DAP [8] to 40 minutes [10]; the use of an in-between value of 20 minutes for full dark adaptation is recommended by Kelbsch et al. (29). Since we planned on using comparatively long experimental light durations of around 15 minutes, we did not want dark adaptation to overly influence sensitivity during the first light steps of the experiment, compared to the last steps, where light adaption would have happened for several minutes, regardless of the DAP. We tried periods of 90 seconds, 180 seconds, and 900 seconds on two participants with the *Up* protocol in three experimental runs, which is described under *Experiment I* below. The longer the DAP, the greater was pupilloconstriction during the first third of the steps (see supplemental *Fig S1*). Compared to a second run of the *Up* protocol directly after the first one, the 90-second DAP pupilloconstriction curve was closest. Therefore, we chose this period for the main experiments.

We also performed preliminary measurements with the *Up* protocol on ten subjects before the main experiments, to explore the behavior of pupilloconstriction in our setup. In those subjects, we found differences dependent on sex, particularly at the longer wavelengths (not shown).

#### Experiment I

*Experiment I* was divided into two protocols, in each of which the stimuli consisted of a stepwise sweep across the available spectrum (*Fig 2*). One such sweep (*Up*) went from the lowest to the highest wavelength, the other (*Down*) from the highest to the lowest. Light onset happened after a dark adaptation period of 90 seconds duration at the beginning of each protocol (see above), and light remained on during the entire protocol. Light steps lasted 15 seconds each, followed by the next light step at a spectral distance of about 5 nm. We hypothesized that with continuous light that shifted slightly in peak wavelength with each step we would see the ipRGC influence in the pupil constriction amplitude by wavelength. Due to the sluggish nature of ipRGC responses [10–12], ipRGC-driven constriction might trail behind, which was an additional reason to balance the spectral protocol series by an up-down alternation. McDougal and Gamlin (11) report that ipRGC-influence at 10 to 18 seconds after light onset takes over as the strongest relative influence on pupil constriction, with rods being a close second; cones were found to have, after 10 seconds, about 1.5 to 2 orders of magnitude lower sensitivity. Since each step in *Experiment I* lasts 15 seconds, we expect a one-step delay to ipRGC pupil constriction, which would equal a 5-nm wavelength shift when looking at just one protocol. Each protocol consisted of 61 light steps, totaling 915 seconds of light (15.25 minutes), or 1005 seconds including dark adaptation (16.75 minutes). Seventy-five participants were enrolled in *Experiment I*. Due to technical difficulties, the *Down* protocol was not available when *Experiment I* started, and was therefore measured consecutively to the *Up* protocol. Due to this sequence, the number of partaking subjects differs between the protocols, with 57 participants in the *Up* and 23 in the *Down* protocol. Of those taking part in the *Down* protocol, we measured five randomly selected participants in both protocols to check whether there were any systematic differences between the protocol groups, which was not the case. Mixed models were used in the statistical analysis to take these factors into account.

**Fig 2.**
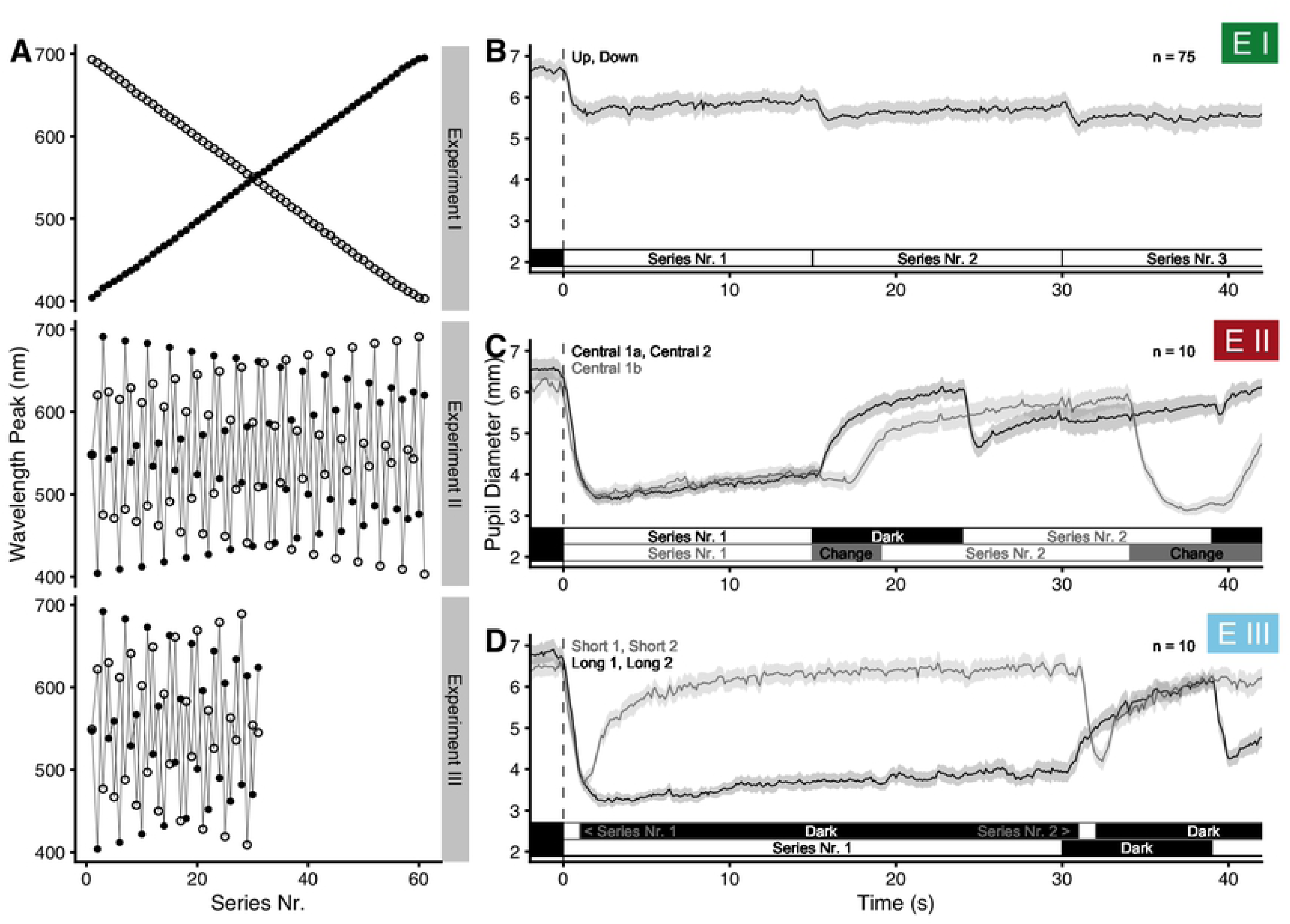
Experimental Protocols. (A) Series of peak wavelengths for each experiment. Filled and empty circles differentiate the series used in an experiment. (B, C, D) Schematics of the first 40 seconds of each experimental protocol. Light onset after dark adaptation occurs at zero seconds (dotted black line). Traces show the average pupil diameter, with a ribbon representing the SEM. For visual reasons, only one of the two protocols sharing the same procedure is shown. These are *Up* (B), *Central 1a* (C), and *Short 1* and *Long 1* (D).

#### Experiment II

The surprise outcome in *Experiment I* was the comparatively high wavelength of 540 nm at which pupil constriction amplitude was at its maximum (see below under *Results* for all outcomes and the corresponding figures). In *Experiment II*, we thus tested whether this was due to our experimental design of small step changes along the spectrum in *Experiment I*. We therefore designed three new protocols (*Fig 2 A* and *2 C*). All three protocols started, after a dark adaptation phase of 90 seconds, at 548 nm, which is halfway on the available range of wavelengths. The protocols *Central 1a* and *Central 2* continued with differing spectral-wavelength shifts between steps. Both comprise 61 light steps in total. Except for the first step, the protocols are symmetrical to the midpoint of the series axis. Since the monochromator required more time for the required large changes of wavelength between steps than in *Experiment I* (in which the change was near instantaneous), the shutter was closed during the wavelength adjustment. In the longest case, the wavelength change lasted just under 9 seconds, and we set the shutter closure time to that value for all steps. The result, for both protocols, was 15 seconds of light followed by 9 seconds of darkness at each step, totaling 24 seconds per wavelength; the total series had 1464 seconds after light onset and 1554 seconds including dark adaptation, i.e. just under 26 minutes). Since the presence of dark periods between light steps introduced a new factor to the experimental design, we designed the further protocol *Central 1b*. This third protocol was identical to *Central 1a*, except that the shutter stayed open at all times, i.e., there were no dark phases in between light steps, and the 15 seconds per light step started right after the wavelength adjustment (*Fig 2 C*). As an undesired side effect, participants perceived the adjustment of wavelengths in that series. We reasoned, however, that these brief periods (about 2.8 ms per 1 nm wavelength) would not influence pupil constriction amplitude 10 to 15 seconds later in a relevant manner.

Ten participants took part in *Experiment II*. The sample size was chosen based on the first experiment. There, we saw that a random sample of ten out of the available sample for every protocol resulted in the same primary outcome for the spectral dependency. Each participant took part in every protocol.

#### Experiment III

Two main results in *Experiment II* led us to the design of a further experiment. *Experiment II* had shown that the spectral dependency of the PCA depended on whether or not there were periods of darkness between light steps, i.e., whether the light stimulus was discontinuous or continuous. Of particular interest was an apparent shift in wavelength sensitivity over time when light steps were discontinuous, but not otherwise. The experiment had further allowed looking at the wavelength dependency of the PIPR during the intervals of darkness. We designed *Experiment III* to build on these results. It comprised four protocols (see *Fig 2 D*), each of which consisted of 31 light steps following the 90-second dark adaptation phase. Light steps followed the series structure of *Central 1a* and *Central 2* from *Experiment II*; however, since each light step was longer in *Experiment III*, we halved the number of steps to not overly tire participants. The spectrum was therefore divided into steps of about 10 nm wavelength difference. The series *Long 1* and *Long 2* were designed to analyze wavelength-dependency changes over a more extended period of time. Accordingly, each step’s period was extended from 15 to 30 seconds duration, followed by 9 seconds of darkness between the steps (1209 seconds after light onset; 1299 seconds including dark adaptation, or about 21.5 minutes). The protocols *Short 1* and Short *2* were designed to analyze the PIPR over a more extended period. Accordingly, each step’s light stimulus lasted 1 second, with 30 seconds of darkness between steps (in total 961 seconds after light onset; 1051 seconds including dark adaptation, or about 17.5 minutes). Ten participants took part in *Experiment III*. Each participant took part in every protocol.

### Procedure

Participants arrived at the appointed time in the laboratory and were welcomed and seated. Measurements were restricted to daytime hours and were scheduled from 08:00 am to 08:00 pm (taking place, on average, at 01:30 pm ± 2:45 h:m). The laboratory was a windowless room with constant temperature, mechanical ventilation, and lit with warm-white artificial light (about 50 to 100 Lux, depending on position). Participants read and signed the prepared informed-consent form. Color vision was tested first. Illumination on the tests was cool white, with a high color-rendering index. Participants then filled out a three-part questionnaire, (1) the German translation of the Morningness-Eveningness Questionnaire for testing chronotype (D-MEQ; scores the time of preference; according to the MEQ manual [24], chronotype scores from 60 and above are considered *morning types* or *Larks*, below 40 *evening types* or *Owls*, and in between *Neutral types*); (2) general demographic questions, and (3) general questions regarding participants’ current health. The experimenter then fitted the pupillometry apparatus to the participants’ head and the participant lied down on a flat treatment couch with a small pillow under the head. The experimenter ensured that the infrared cameras were positioned correctly so that pupil diameter could later be calculated from camera pictures, either by adjusting the distance to the eye (in *Experiment I)* or by presenting a reference scale at eye level (in *Experiment II* and *III*). The Ganzfeld dome was then pivoted over the participants head until the eyes were at a predetermined position and only the dome’s inside was visible. Room lighting was then switched off, and a black, opaque curtain between the participant and the experimenter was drawn shut. Thereby the eye-tracker camera output could be monitored on a personal computer without stray light influencing the experimental setup. Participants were instructed to relax and look straight ahead, with minimal blinking. The experimenter repeated the instructions in following procedures if compliance faded. The appropriate protocol was then started, each beginning with its 90-second dark adaptation phase, followed by the respective series (see above). Participants’ eyes were not medically dilated, and both eyes received the light stimulus (closed-loop paradigm [29]). If participants took part in more than one protocol (primarily in *Experiment II* and *III*, see above), protocol order was randomized. Participants further got up from the treatment couch for about five minutes in-between protocols during which room lighting was switched on, as in the initial setting. At the end of the experimental session, participants were thanked and debriefed. The total time participants spent on site was between thirty minutes in *Experiment I* and about two hours in *Experiment II* and *III*.

### Data analysis

Bio-signal data analysis covered converting raw pupil data to a time series of constriction amplitudes for each participant and protocol. It further covered spectral calculations to derive photopic, mesopic, and scotopic illuminance values for each monochromatic wavelength peak.

#### Pupillary constriction amplitude

We used the *R* software (Version 4.0.2) [30] with several packages for data analysis (*anytime*, *cowplot*, *ggplot2*, *ggmisc*, *knitr*, *readxl*, *signal, tidyverse)*. Raw pupil data were stored in pixel-based units by the measurement software, and exported to CSV files. These values were converted to mm and mm^2^ by a factor derived from the *ImageJ* software [31], using screenshots from the eye tracker videos with included reference scales (see above). Blinks and other artefacts were then removed by calculating a *circularity index* (ratio of pupil height to width, and width to height). Pupil values with a *circularity index* smaller than 0.7 were removed [10, 32]. We also removed pupil diameter values outside reasonable thresholds, usually when above 8.5 mm or below 1.5 mm pupil diameter. However, in some cases, a different cut-off was decided upon after visual inspection. A Savitzky-Golay filter [33] with a third-degree polynomial [22, 34] was used for smoothing of the pupil data; filter length was set to 31 data points or about 0.5 seconds. The same filter was also used to calculate the second derivative of the smoothed curve – valleys in the second derivative coincide with the early stages of pupil constriction. The timestamps of these valleys were used to shift the measurement time according to light onset. The shift was monitored visually and, when necessary, manually adjusted. After this correction, time values are negative during dark adaptation, light onset happens at zero seconds, and the experimental lighting conditions occur at positive time values according to the protocol.

In the next step, the pupillary constriction amplitude (PCA) was calculated at each time step *i, as*

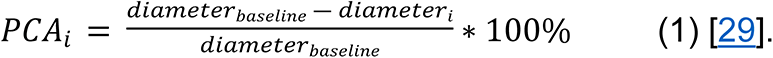

Pupillary constriction amplitude is the decrease of pupil diameter, normalized to a baseline level (generally the baseline before stimulation [29]), thereby controlling for the considerable variation of inter-individual pupil size, as well as for its (smaller) intra-individual variation [14, 29]. Baseline diameter corresponds to 0% constriction, and the (impossible) pupil diameter of 0 mm is considered 100% constriction. As an example, a light-adapted diameter of 2 mm from a baseline pupil diameter of 8 mm equals 75% constriction amplitude. As we used a comparatively short dark adaptation period, followed by a long series of light steps, we believe that the pupil diameter at the end of the dark adaption phase somewhat loses its prominence, as it is mostly unrelated to any particular wavelength peak across the protocols. Therefore, we used the largest one-second-mean pupil diameter (see below) as the pupil baseline, taken from the period between the last seconds of dark adaptation until the end of each protocol. This baseline still takes interindividual differences into account, while intraindividual differences are already taken care of through the continuous measurement over all wavelengths per person and protocol. By definition, PCA can only take positive values here. To check whether our normalization method affected the results in an undesirable way, we performed an additional primary analysis without normalization in *Experiment I*, and further tried an alternative normalization in *Experiment II* and *III* to the average diameter during the last second of darkness before each light step. Since we had measurements for both eyes, one eye was discarded at the next step; the eye with the highest count of remaining data points was kept unless visual inspection showed abnormalities. The CSV export from the collected steps above contained the pupil diameter (mm), PCA (%), Series number, time since dark adaptation/light onset (s), time since the start of the light step (s), wavelength (nm), baseline diameter (mm), participant code, eye, date, time, and protocol name. Data in the CSV export file were further aggregated in two ways, so besides unaggregated pupil data (i.e., with 60 Hz sampling frequency) the file contained one-second means, and means over specific periods. The specific periods were the last five seconds of each light step, for every protocol except *Short 1 or Short 2,* in which the sixth second after light offset was used instead [17]. In a final step before the statistical analysis, the pupil data were combined with participant data (sex, age, chronotype), and spectral data (see *Fig 1* and below). The scripts to derive the input data for the statistical analysis are available as *R-Markdown* files as part of the *Supporting Information*, in the S*2 file*. All data used in the statistical analyses are available as CSV files from the *Open Science Framework* [35].

#### Spectral calculations

From the spectral irradiance measurements, total irradiance was calculated automatically by the measurement software (see above). Similarly, illuminance values were automatically calculated according to the CIE-1931 standard 2° photopic observer which uses the V(λ) function [36], as well as according to the CIE-1964 standard 10° photopic observer, which uses the V_10_(λ) function [37]. The five types of alpha-opic equivalent (D65) daylight illuminance were calculated according to the CIE S 026 standard *(CIE-S-026-EDI-Toolbox-vE1.051*) [28]. We also used the *Melanopic-light-sources_Toolkit_V13.12*, which was developed and provided to us by Dieter Lang [38]. The toolkit provided illuminance values for the CIE-1951 scotopic observer with V’(λ) [39] spectral weighing in addition to the alpha-opic values. Illuminance for the mesopic observer was calculated according to the German standards DIN 5031-2 [40] and DIN 5031-3 [41] through V_eq_(λ) weighing functions. The standards were recommended by Adrian (5). The CIE has since released the standard CIE 191:2010 for mesopic photometry, which is based on the same basic principles but uses a different calculation procedure and notation (V_mes;m_(λ)) [42]. In the German standards, illuminance calculations for the mesopic observer use spectral weighing functions V_eq_(λ) that are intermediary between the photopic V_10_(λ) and the scotopic V’(λ) function, in addition to intermediary luminous efficacies of radiation. These intermediaries are chosen based on the adapting equivalent luminance L_eq_ for a 10° observer on a log_10_-based scale between the photopic (100 cd/m^2^) and scotopic (10^-5^ cd/m^2^) endpoints defined by the standards. Our calculations were performed using *Microsoft Excel* software. The spectral measurements for the range of peak wavelengths, including total irradiance and all types of illuminance values are provided as part of the *Supporting Information*, in the *S3 file*.

#### Statistical methods

We used the *R* software (Version 4.0.2) [30] with the *mgcv* (Version 1.8-31) [43] package to perform a *generalized additive mixed-effect analysis* on the connection between pupillary constriction amplitude and the empirically and theoretically derived predictors. *Generalized Additive Models* (GAMs) [43, 44] allow for a data-driven decomposition of the relationship between a dependent variable and user-defined predictor variables in both a parametric and nonparametric fashion. A variant of GAMs are *Generalized Additive Mixed-Models* (GAMMs or HGAMs), used in the context of hierarchical data as is the case in any repeated-measures setup such as ours [45]. GAMs are widely used in *Biology*, *Ecology*, and *Linguistics* [43]. GAM(M)s are not yet used often in *Human Life Sciences* [46], but we find them an excellent match for the analysis at hand, and mixed-effect models gain support in biophysiological research such as visual perception [47] and nonvisual effects of light [48]. Guidelines on the theory behind GAMs and their practical use can be found in Wood (43), Simpson (49), Pedersen et al. (50), and the guide by Sóskuthy (45).

The nonparametric parts of GAMs take the form of *smooth* functions (*smooths*, in short), describing the connection between the outcome variable and a predictor. A *smooth* function *f*(*x*_*i*_) is the weighted sum of a number *k* of basis functions *b*_*j*_ at the *i*-th value of a predictor variable *x*:

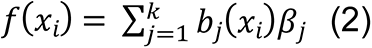

The *β*_*j*_ in *eq 2* denote the weights (the *smooth* coefficients) of the basis functions *b*_*j*_(*x*_*i*_). Since GAMs are not yet widely familiar in this field of research, their concept is visualized in *Fig 3 A*. The type of basis function is chosen as part of the model construction, with *cubic regression splines* being probably the most well-known type (*Fig 3 A1*). Unless otherwise stated, we used the *mgcv*-default type of basis functions, which are *thin plate regression splines* (TPS). Wood (43) gives a comprehensive overview of basis functions and their suitability depending on the context. Unlike coefficients in a normal parametric regression, the basis-function weights *β*_*j*_ in generalized additive models are not by themselves interpretable towards the modelled relationship (*Fig 3 A2*). Only by knowledge of the set of basis functions and their eponymous weighted addition (“*additive” models*) can the resulting *smooth* function be interpreted (*Fig 3 A3*). One of the underlying assumptions to the *smooths* is a constancy of complexity across the value range of *x*. That means that *smooths* do not do well without additional adjustments, if constructed from relationships that vary heavily in their dynamics across the range, or that exhibit discontinuous changes (step-changes). These adjustments include the use of *adaptive splines* for varying dynamics, or the use of factors for step-changes [43]. *Smooth* functions can be multi-dimensional, thereby describing interdependent relationships or interaction effects between predictors and the outcome variable. The number *k* of basis functions is chosen based on the complexity of the modelled relationship present in the data. Overfitting is avoided through the penalization of complexity as part of the likelihood optimization. Therefore, added complexity (or *wiggliness* [43]) in the *smooth* needs to improve the model fit enough – compared to, e.g., a straight line – to outweigh the penalization. More broadly, GAMs ascertain to model the simplest possible relationship between variables, or no relationship at all, without leaving out relevant structural components depending on any given predictor.

**Fig 3.**
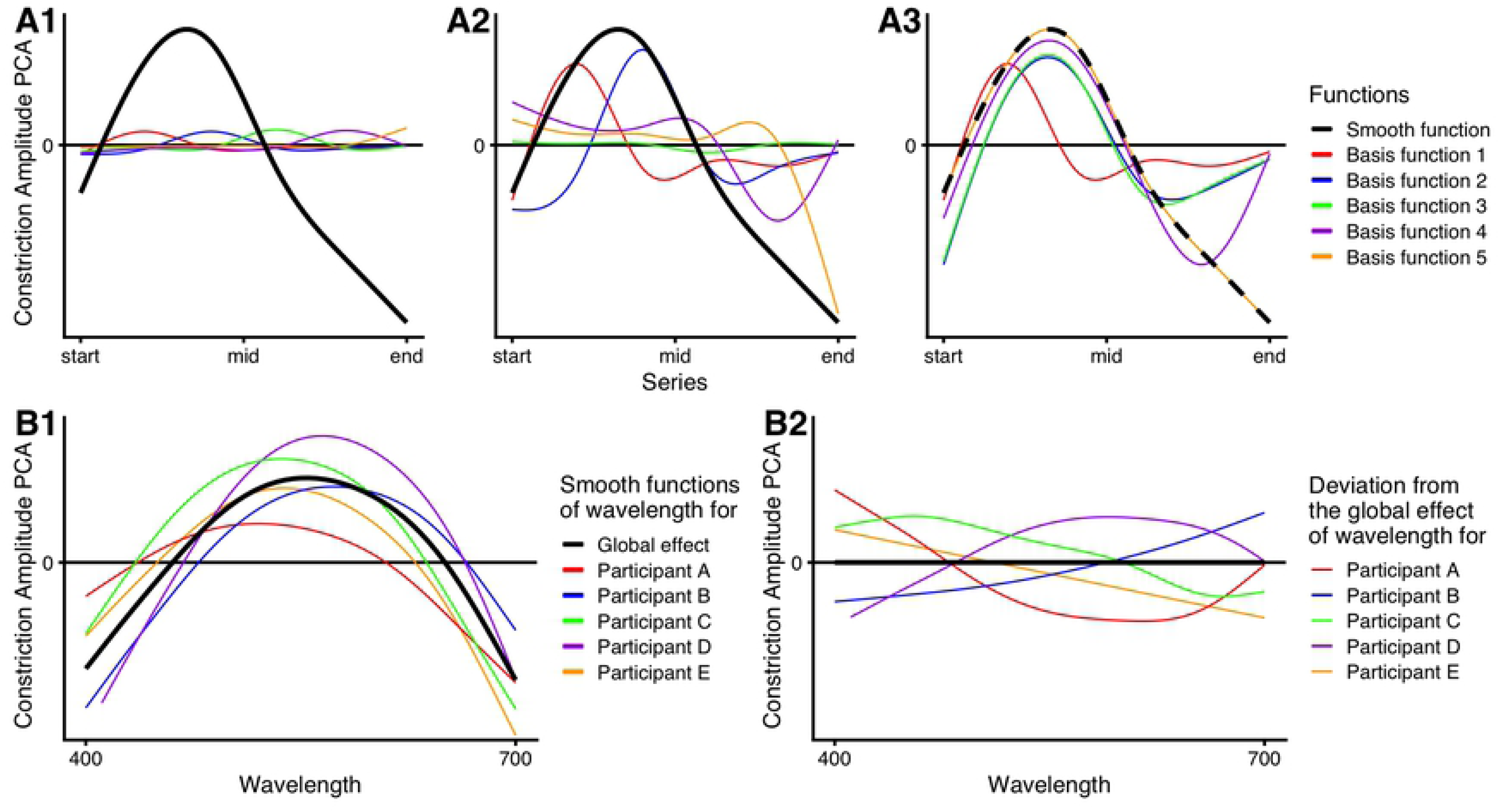
Concepts of additive (mixed-effect) models. (A) Construction of a *smooth* through *basis expansion*. Every *smooth* is constructed from a number of basis functions, usually spread evenly across the value range of a predictor. In the shown case, five cubic regression splines were used as basis functions to demonstrate the concept. The thick black line represents the final *smooth* function, describing the relationship between the predictor and the outcome variable. Colored traces show the basis functions. (A1) Unscaled basis functions – their type and maximum number are part of the input for model generation. (A2) Scaled basis functions – for each basis function, a respective weight is estimated by which the function is scaled. (A3) Summation of basis functions, starting with the first and then, successively, adding the others; the generated *smooth* can then be used for prediction. The *smooth* is shown as dashed curve to show that, with the addition of the last basis function (orange line), the resulting curve is equal to the *smooth* function. (B) Concept of global effects with random *smooths*. (B1) The thick black line represents the global effect, describing the average relationship between the predictor and outcome variable. Colored traces show the individual’s effect of the same predictor, demonstrating interindividual differences. The model takes these differences into account through the so-called random *smooths* (i.e. “random” in that their contribution depends on the subject). (B2) Random *smooths* are *smooths* describing the deviations from the global effect. The colored traces show the deviations present in panel B1. Because the global *smooth* and the random *smooths* are estimated together, the deviations disappear on average, i.e., not all deviations will tend in the same direction, but are spread around the global effect.

If not stated otherwise, we used *Akaike’s Information Criterion* (AIC) for model selection, as suggested by several sources on GAM(M)s [43, 45, 49–51]. The AIC incorporates model fit (likelihood) and model size (number of parameters) [43]. It can further be compared between similar models, with the lower value indicating the better-suited model. Following the example of Pedersen et al. (50), models that differ by two units or less from the lower AIC have substantial support, with the more parsimonious model to be preferred. We report ΔAIC values when there is support for the inclusion of a predictor. T-tests obtain P-values for parametric terms. The p-values for nonparametric terms are *approximate p-values*, so called owing to the complexity of the *degrees-of-freedom* (*df*) concept used for statistical testing, and other underlying assumptions [43, 45]. For nonparametric terms, the hypothesis tested-against states that the relationship between predictor and dependent variable is a horizontal, flat line. P-values less than or equal to 0.05 were considered significant. Confidence intervals in prediction plots account for the model’s overall uncertainty, not for the respective plotted predictor alone.

We explored the PCA relationship with *wavelength*, *irradiance*, *series, time*, *time of day*, *sex*, *chronotype*, and *age* by the methods described above. We further looked for the PCA’s best fit to *photopic, scotopic, and mesopic illuminance*, and *alpha-opic equivalent-daylight illuminance* values. The variable *wavelength* refers to the dominant or peak wavelength of the monochromatic stimuli as described above. After light offset, *wavelength* refers to the prior light stimulus. *Series* refers to the numbers of light steps after the dark adaptation phase. *Time* refers to the time since the start of the current light stimulus, i.e., light onset or change in wavelength. Several additional R packages were used for the statistical analysis and plot generation (*tidyverse*, *ggplot2*, *readxl*, *dplyr*, *cowplot*, *lubridate*, *itsadug*, *printr*, *patchwork*, *here, reshape2, plotly*, *gganimate*, *gridGraphics*, *transformr*, *glue, magick,* and *DT)*. All scripts for analysis and plot generation are available as *R Markdown* files in the *Supporting Information*, S*4 file*.

#### Base model structure

While the relevant code is included as part of the *S4* file, we believe it helps explaining the basic model structure and settings used to analyze the experimental outcome. The model structure is one of five basic GAMM variants (according to Pedersen et al. (50)), which differ in how fixed and random effects are included. The chosen variant, by way of AIC comparison and model diagnostics, is shown in the following equation:

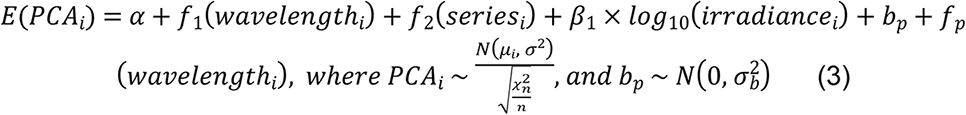

*Eq 3* shows the statistical base model, where the expected value of PCA *E*(*PCA*_*i*_) is modelled by *smooths* as a function of *wavelength (f*_1_) and *series(f*_2_), and a parametric effect of *log_10_(irradiance)*; *α* is the average PCA, when all other terms are zero, also called the intercept; *β*_1_ is the parametric coefficient, or slope, for *log_10_(irradiance)*; *b*_*p*_ are random intercepts by participant; and *f*_*p*_(*wavelength*_*i*_) are random *smooths* by participant.

*Wavelength* is the main predictor, *series* was included for empirical reasons, i.e., in the cases when visualization of the PCA data in *Experiment I* showed the influence of the protocol (*series)*. As stated above under *Apparatus*, *irradiance* is theoretically motivated to compensate for lower irradiance levels under 450 nm wavelength. However, *smooth* behavior from 400 to 450 nm still has to be interpreted tentatively, since the estimate for *irradiance* might under- or overcompensate the effect, thereby distorting the influence of the lowest wavelengths on PCA. We account for this by leaving *irradiance* out of the model in a secondary analysis and discuss differences between the variants.

As random effects in *Experiment I* we had random intercepts *b*_*p*_ and random *smooths f*_*p*_ (*wavelength*_*i*_) for each participant *p*. In *Experiment II* and *III*, all participants took part in all protocols; we, therefore, had random effects for participants-by-protocol in those cases [45]. Random intercepts describe the deviation from the average PCA (*α*) depending on the participant (or participant-and-protocol), with independent and gaussian distributed values. Random *smooths* can be thought of as the nonparametric version of random slopes in *linear mixed-effect models*. Random *smooths* allow for the individual’s deviation from the global effect *f*_1_(*wavelength*_*i*_), but zero-out on the global effect when viewed on average across participants (*Fig 3 B*).

For the distribution of our response variable, *PCA*_*i*_, which takes positive values between 0 and 1, the *gamma* or *beta* distribution seemed sensible assumptions initially. However, model diagnostics with those were poor compared to even the standard *gaussian* error distribution. Instead, we thus went with the *scaled-t* (*scat in mgcv*) distribution family, which is less susceptible to outliers and indeed greatly improved residual distribution. Eq 3 states that *PCA*_*i*_ varies around its mean *μ*_*i*_ with a t-distribution based on the standard deviation *σ* and *n* degrees of freedom. We also tested a model with the *gaussian location-scale* (*gaulss in mgcv*) family. This distribution allows the variance to change depending on a predictor, just like the mean. However, we did not find relevant differences in variance, and the model with the *scat* family performed similarly well with respect to the AIC. The *gaulss* family did further not allow for the computationally faster *bam* command, so we used the *scat* family for the analysis.

We used the *gam.check* function of *mgcv*, which includes several residual plots for model diagnosis. Visual inspection of these plots did not reveal any apparent deviation from homoscedasticity or normality. Models were further controlled for autocorrelation in the residuals. In all models that used only one timestep per wavelength peak (5-second average PCA), the inclusion of random effects diminished all relevant autocorrelation. In models that used 1-second averages for PCA, random *smooths* by *wavelength* left serious residual autocorrelation present. Random *smooths* by both *wavelength* and *time* proved too computationally intensive in those cases and we instead included the remaining autocorrelation with an autoregressive error model [45, 49], in addition to random *smooths* by *wavelength*. Finally, the number of knots for each *smooth* (i.e., how many basis functions comprise the *smooth*) was also checked (*k.check*) and was increased, when theoretically sensible, until autocorrelation along the *smooth* vanished [50]. For computational speed reasons, we used the *bam* function with *fREML* in R instead of the standard function of *gam* with *REML*. *Bam* is optimized for big data sets and model structures.

If not otherwise stated, GAMs in the *Results* section were used to model pupillary constriction amplitude (PCA) according to *eq 3,* the latter also called the base model. Additional parametric and nonparametric predictors were explored – their addition to or removal from *eq 3* are stated in the respective sections. The default PCA is the average value of each light step’s last five seconds, as described above.

#### Linear mixed-effect model structure

We used the *lme4* package [52] to perform a *linear mixed-effects analysis* of the relationship between PCA and the various illuminance variables described above:

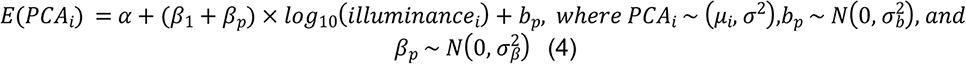

Intercept *α* and slope *β*_1_ are the fixed effects of the model described in *eq 4.* The intercept *α* indicates the value of PCA, when all other terms in the equation are zero. The slopes, or beta coefficients, *β*_1_ and *β*_*p*_, represent the change in the expected value of *PCA*_*i*_ when increasing illuminance by one log unit. As random effects we included random intercepts by participant, *b*_*p*_, and random slopes by participant, *β*_*p*_, both of which have independent values with a *gaussian* distribution. Random intercepts *b_p_* show how much individuals deviate from the average PCA *α* with a variance of 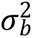. Random slopes, similarly, show how much the relationship between the predictor and outcome variable deviate from the fixed effect *β*_1_ with the variance 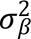. *PCA*_*i*_ has a *gaussian* distribution, with variance *σ*^2^ around its mean *μ*_*i*_. The proportion of variance-accounted-for (R^2^) was calculated according to Xu (53).

## Results

An overview of the experimental data and circumstantial conditions is shown in *Fig 4*. *Fig 4 A* shows plots for PCA vs. wavelength for each protocol, where the underlying data are not controlled for any dependencies. *Fig 4 B* shows *chronotype* vs. *time of day*. Noteworthy is the lack of *Owl* chronotypes before midday, as well as pronounced *Larks* in the morning. *Fig 4 C* shows at what time in the year the experiments took place, as suggested by Veitch and Knoop (54). Experiments took place in 2019 (preliminary measurements, *Experiment I*) and in 2020 (all experiments).

**Fig 4.**
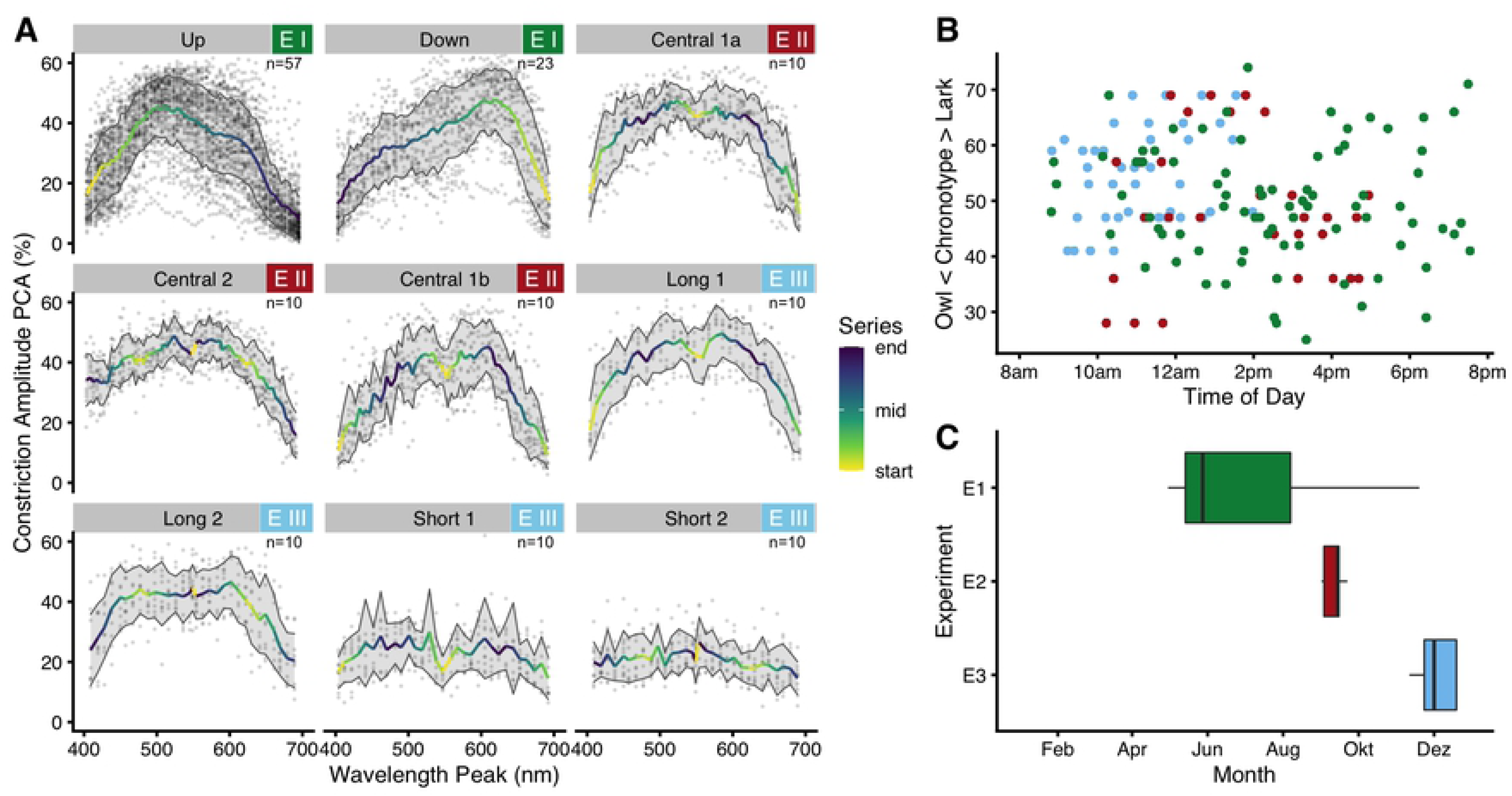
Experimental data and circumstances. (A) Pupillary constriction amplitude (PCA) plotted against wavelength, for each of the nine protocols. The color scale shows at which point in the series a specific wavelength was presented; light yellow represents early in the series. For all but the *Short* protocols, points represent the average PCA during the respective last five seconds of a light step. In the two *Short* protocols, points represent the average PCA during the sixth second after lights-off (or sevenths second after lights-on). Traces show the mean PCA, ribbons its standard deviation. The number in the upper right corner of each plot shows the corresponding sample size. (B) Scatterplot of when subjects of a certain chronotype started their respective protocols. The color scheme is according to (C), i.e. green, red, and blue correspond to Exp. I – III, respectively. Note the lack of *Lark* and *Owl* chronotypes in *Experiment I* before midday. (C) Boxplot of the time of the year when the experiments took place. Preliminary measurements and parts of *Experiment I* took place in 2019, the other measurements in 2020.

### Experiment I

#### Base model results

The results for the base model of all three experiments are shown together in *Fig 5,* for better comparison. The results for *Experiment I* are shown in *Fig 5 A* and are reported in this section; the results for *Experiment II* and *III*, as well as secondary results for *Experiment I* are shown here in *Fig 5 B*, *C*, and *D*, respectively, but are reported below in the respective sections. The statistical base model contains *wavelength* and *series* as nonparametric predictors, and *log_10_(irradiance)* for the average PCA during the last five seconds of each light step as a parametric predictor. There is strong support for a dependence on *wavelength* in its implemented form versus other random effect structures (minimal ΔAIC = 33), or no wavelength dependence (ΔAIC = 4963), as there is for including *series* as predictor (ΔAIC = 390). *Irradiance* did improve the model further (ΔAIC = 17). All p-values in the final model were below 0.001. The dependence on *wavelength* shows an inverted-U shaped curve with a peak at 540 nm, and an average PCA of about 38% when controlling for all other factors (*Fig 5 A1*). Compared to the maximum, PCA is predicted by the model as up to 11% lower for short wavelengths and up to 17% lower for long wavelengths. The dependence on *wavelength* shows a peak at 550 nm when not controlling for *irradiance* (supplemental *Fig S15A*). With respect to series effects, PCA increases slightly with *series* up to step 18 (by about 3%), which occurs at about 360 seconds after the protocol start, or, respectively, 270 seconds after lights-on (*Fig 5 A2*). After this point, PCA declines by about 13% until the end of the protocol. PCA further increases with *irradiance* (β_log10(irradiance)_ = +6.5% ±1.4% SE), as seen in *Fig 5 A3*. 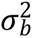 was 8% in *Experiment I*, which is the standard deviation of the random intercept by participant.

**Fig 5.**
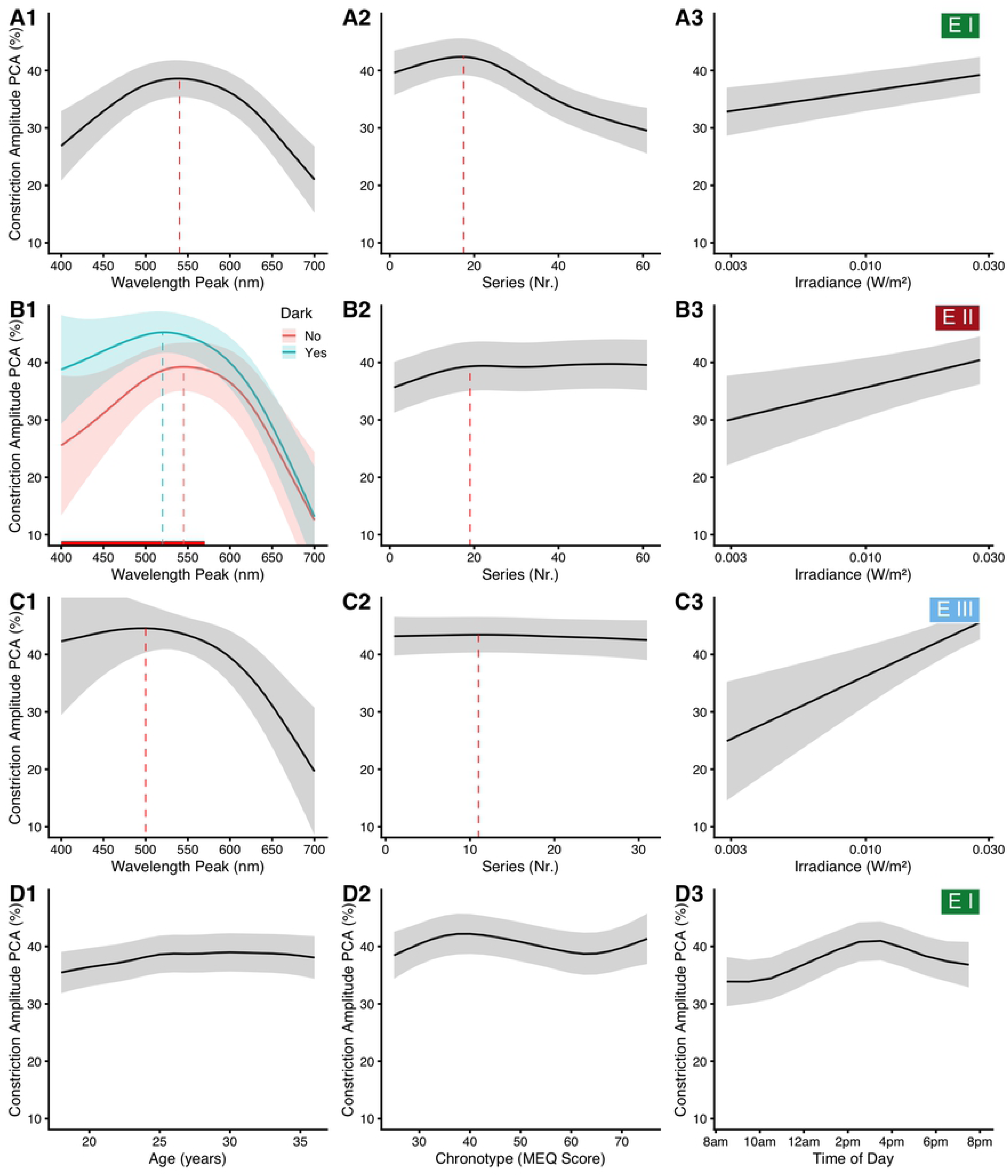
Model predictions. Model predictions for pupillary constriction amplitude (PCA) as depending on several main predictors, when all other predictors are held at an average, constant level. Traces show the model prediction for the mean, ribbons its 95% confidence interval. Dotted lines show a peak that is discussed in the main text. In (A3, B3, and C3), the x-axis is logarithmically scaled to reflect the logarithmic transformation of *irradiance* in the model. (A) and (D) show dependencies in *Experiment I*, discussed here, (B) and (C) dependencies in *Experiment II* and III, respectively, discussed later. (B) The red line at the bottom of (B1) indicates where the two curves differ significantly at the 0.05 level of significance.

#### Interaction of series and wavelength

There is some support for an interaction of *wavelength* with *series*, i.e., that changes of PCA with *wavelength* depend on when, in the series, specific wavelengths are presented (ΔAIC = 5). Results for that are shown in a supplemental figure (S5). The model prediction with the interaction effect shows little or no deviation from the basic model results at the instances with data. Compared to the model without interaction, PCA is predicted to be lower for very short and long wavelengths occurring right after light onset. Wavelengths around 600 nm further lead to a more pronounced PCA during the early steps of the series, while for wavelengths around 500 nm PCA is slightly lower during the later steps. In terms of ipRGC sensitivity, it seems as though, compared to the model without the interaction, PCA is heightened at, and a few steps after, reaching the sensitive wavelengths around 490 nm, but the effect is small. With the interaction effect as part of the model, *irradiance* is no longer significant (p = 0.31).

#### Sex and age

There is further support for an influence of *sex* on the PCA (ΔAIC = 85). On average, women had a smaller PCA than men (β_Women_ = –4.4% ±0.4% SE, p < 0.001). Our preliminary measurements indicated that the *wavelength* dependency of PCA is conditional on *sex* (not shown). There was, however, no interaction of *sex* with either *wavelength* (p = 0.98) or *series* (p = 0.27) based on the larger sample from *Experiment I.* With *age* as a predictor (*Fig 5 D1*), PCA increased by about 3% from age 18 up to about age 25 (ΔAIC = 245, p < 0.001). At higher age, PCA reached a plateau. We did not see changes in *wavelength* with *age* (p = 0.67).

#### Chronotype and time of day

There is strong support to include *chronotype* and *time of day* as predictors, together with a three-way interaction of these with *wavelength* (ΔAIC = 47 and ΔAIC = 130, respectively). Main effects of *chronotype* and *time of day* are shown above in *Fig 5 D2* and *D3.* In general, *Owls* have a slightly higher PCA compared to *Larks* (*Fig 5 D2*), and PCA was highest during the early afternoon around 3 pm (*Fig 5 D3*). According to the interaction model, subjects with neutral chronotype (chronotype score of 50) showed no substantial change in wavelength dependency across the day (*Fig 6 A2,* blue curves). In contrast, subjects with pronounced chronotype did (*Fig 6 A2,* green and red curves; for a complete overview from 8:30 am to 7:30 pm see supplemental *Fig S7*). *Larks* (green traces) were shifted in sensitivity towards short wavelengths before midday, were about equal to *Neutral* types at around noon and early afternoon, and were more sensitive to longer wavelengths in the late afternoon. *Owls* (red traces), on the other hand, were shifted opposite; before midday they were more sensitive to longer wavelengths, were close to *Neutral* types at midday and early afternoon, and were more sensitive towards shorter wavelengths during the evening hours. Not every point of this three-dimensional interaction structure has enough data points to support all predictor variable combinations. Missing combinations occur in particular for extreme chronotypes and measurement hours, as was shown in *Fig 4 B* above. However, enough structural points are present to support the dependency thus described (see supplemental *Fig S8* for more details). We pooled data from all experiments to increase the number of combinations. The results from the pooled data model are shown in *Fig 6 B*, but are reported further below.

**Fig 6.**
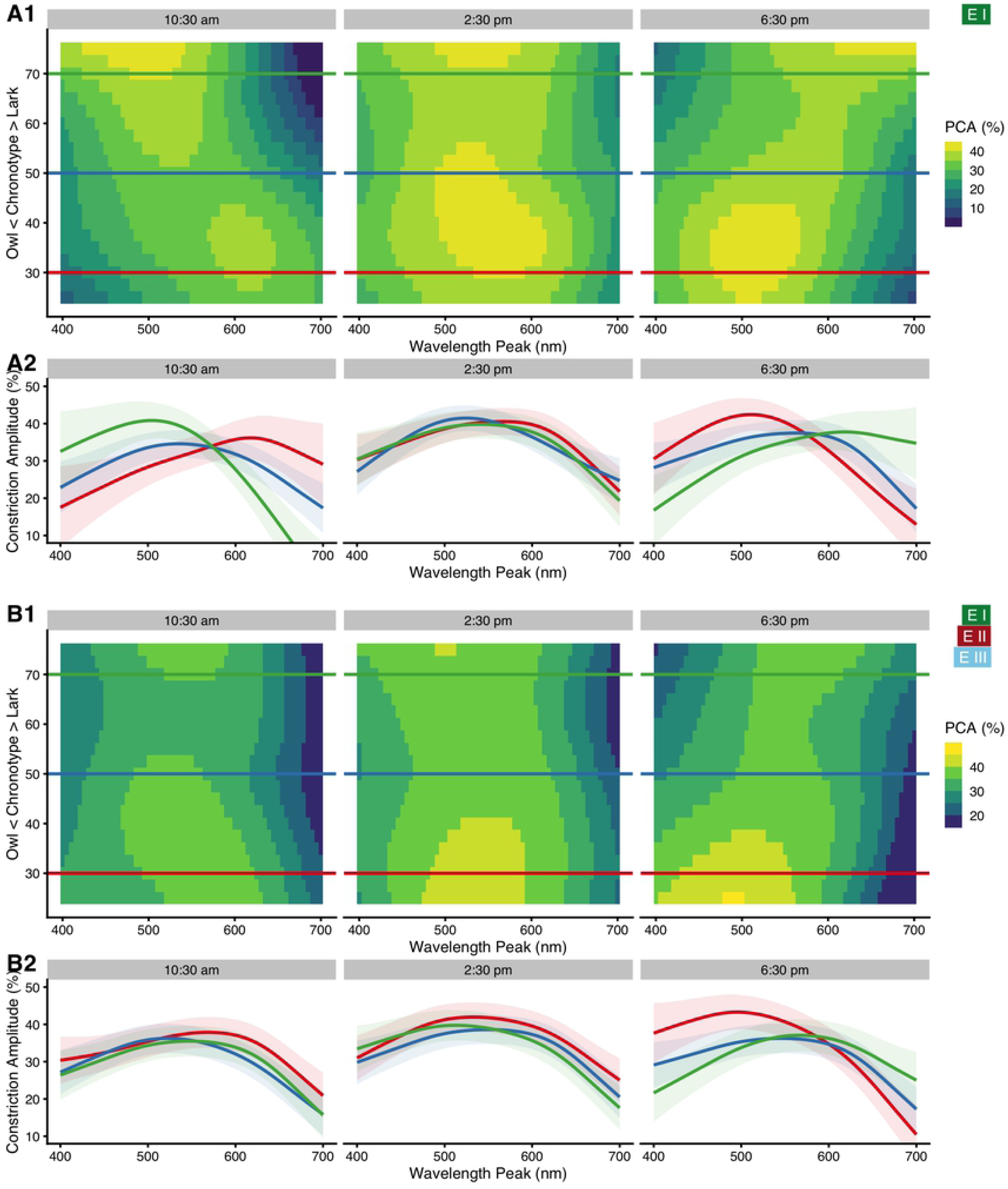
Interaction of *wavelength*, *chronotype*, and *time of day*. (A1) False-color graph of model predictions for the PCA’s dependence on *wavelength* (x-axis) and *chronotype* (y-axis) for three times of day, when all other predictors (basic model) are held constant at their average. Horizontal lines show where the respective three traces shown in figure part (A2) are taken from. (A2) Model predictions for the PCA vs. wavelength for three chronotypes (trace color), and three times of day. Green traces show *Larks* (CT score = 70), red traces *Owls* (CT score = 30), and blue traces *Neutral* types (CT score = 50). Ribbons show the 95% confidence interval for the predicted means. (B1/B2) Like (A), but for pooled data across all experiments. See the main text for further details.

#### Other Results

When all significant terms are added to the model – i.e., *the interaction of wavelength and series*, the main effects of *sex* and *age,* and further the *interaction* of *chronotype* with *time of day* and *wavelength* – all model terms remain significant (all p < 0.001) except for *irradiance* (p = 0.31). More importantly, the predictor-response relationship across the terms is very similar to the descriptions above. One exception is the effect of *age*. When controlling for *chronotype* and *time of day*, PCA is predicted to gradually rise across the whole range from age 18 to 39 (supplemental *Fig S6 B*), as opposed to only the range from age 18 to 25 (supplemental *Fig S6 A*). This full model was also the best in terms of AIC yet (min ΔAIC = 58). AIC would have been further improved by dropping the main effects of *chronotype* (ΔAIC = 7) or *time of day* (ΔAIC = 40). However, it is not advisable to drop the main effects in the presence of an interaction [43]. Curiously, most of the random *wavelength*-by-participant *smooths* were not significant in this model (61 out of 75). This indicates that the fixed effects above account sufficiently for the interindividual differences in *wavelength* dependency from roughly 80% of participants. PCA was calculated for the centered and scaled pupil size, as described above in *Materials and methods* (eq. 1). When the pupil diameter is not scaled but is centered on the baseline, results show the pupillary constriction in mm instead of in per cent. When pupil diameter is neither scaled nor centered, results show the raw pupil diameter in mm. We explored those two variants of the response variable, but results do not indicate that such scaling changes the model’s composition or interpretation (supplemental *Fig S9*).

#### Illuminance

To analyze how well each type of illuminance can predict PCA, we constructed several simplified standard linear mixed-effect models according to *eq 4,* with a fixed effect for illuminance, and random intercepts and slopes per participant. Illuminance values were log_10_ transformed. Nine measures of illuminance were used as described above in *Materials and methods*: photopic illuminance for foveal and ganzfeld stimulation (2° and 10° observer according to *CIE* standards), mesopic and scotopic illuminance, and the *alpha-opically weighted equivalent daylight illuminances* for the five types of photoreceptor. Since the scotopic illuminance and rhodopic equivalent daylight illuminance use the same underlying action spectrum, results on model fitting are identical between these two types of illuminance. Therefore, only results for the scotopic model will be shown in the figures. *Fig 7* shows the results for all three experiments together for better comparison. The results for *Experiment I* (*Fig 7 A*) show that the photopic (2° and 10°) and especially mesopic functions are much better at predicting PCA (R^2^_illu_ ≥ 0.41) than the commonly used melanopic and rhodopic (viz. scotopic) receptor functions (R^2^_illu_ ≤ 0.24). Of the three cone functions, erythropic and especially chloropic weighing led to good results (R^2^_illu_ ≥ 0.42), whereas cyanopic weighing was the only illuminance type with no support at all by the model (p = 0.37, R^2^_illu_< 0.01).

**Fig 7.**
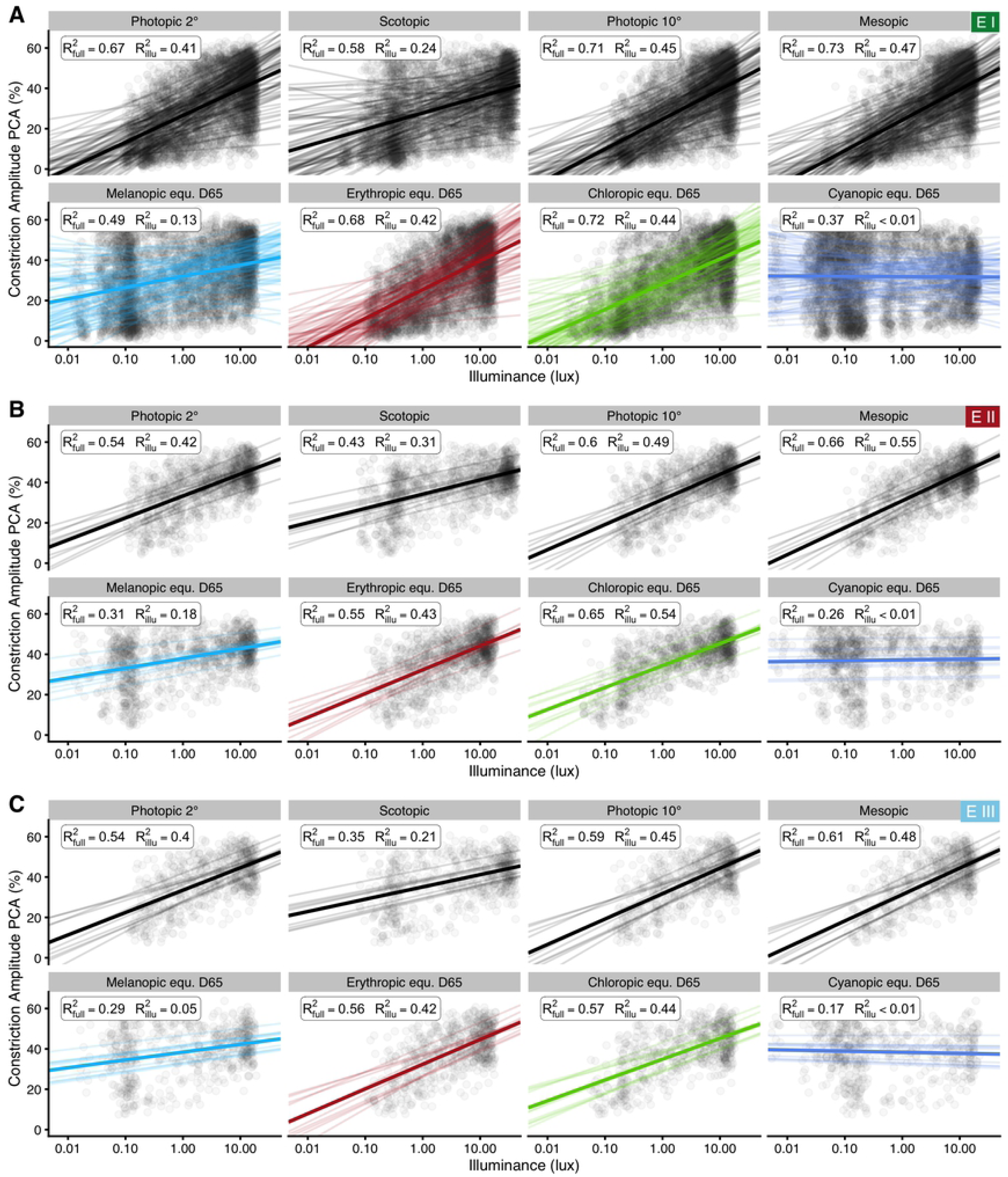
Linear mixed-effect model results for PCA’s dependency on illuminance. Model predictions for PCA vs. various measures of illuminance. Points show individual data. Thick regression lines show the fixed-effect relationship and thin regression lines random effect variation in slope and intercept by participant. The insets show R²_full_ for the full model (fixed and random effects), and, more importantly, a partial R²_illu_, i.e. the proportion of variance explained through PCA’s relationship with the fixed effect of illuminance. Part (A) shows results for *Experiment I*, (B) for protocols with darkness between light steps in *Experiment II*, and (C) for protocols with thirty seconds of light, followed by nine seconds of darkness in *Experiment III*.

Results for Experiment II and III are shown in *Fig 7 B* and *7 C*, respectively, for better visual comparison across the experiments. Their results are reported below in the respective sections.

### Experiment II

#### Base model results

The results of *Experiment II* for the base model were shown above in *Fig 5 B* but are described here. The base model is constructed identically to that in *Experiment I*, with an added factorial predictor for whether there is, or is not, a period of darkness between light steps (referred to as *Dark* in the remainder). There is support for including this factor (ΔAIC = 5). The dependence on *wavelength* shows an inverse-U shaped curve in both cases (*Fig 5 B1*). The difference between the two curves is about 13% PCA in the short-wavelength spectrum but almost disappears towards the long wavelengths, above about 600 nm (it is significant below 570 nm at the 5% level). PCA increases slightly with *series* up to step 19 (about 3%) and is steady afterwards (*Fig 5 B2*). PCA further increases with *irradiance* (β_log10(irradiance)_ = +10.7% ±3.4 SE, *Fig 5 B3*). When excluding *irradiance* as a predictor, the shorter wavelengths become less influential; the difference between the curves remains similar (supplemental *Fig S15B*). 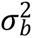 was 5.3% in *Experiment II*, which is the standard deviation of the random intercept by participant.

#### Time-course of the wavelength dependency

As stated above in *Materials and methods*, we aggregated the 60Hz-resolution data to mean values over specific periods, i.e., the last five seconds of light, but also created mean values over each one-second period. We used these one-second values to explore how PCA vs. *wavelength* changed over the time course after light onset for each wavelength (referred to as *time*). The results are shown in *Fig 8 A1* and *A2* (shown in full in the supplemental *Figs S10* and *S11*). Note that this implies an oversimplification for the first one or two seconds, where great changes in pupil diameter occur very quickly (cf. *Fig 2 B*, *C*, and *D*); the purpose here is to get an overview for the later stages of every light step. PCA decreased over the time course (vertical direction in *Fig 8 A1*, horizontal sequence of graphs in *Fig 8 A2*). Short wavelengths are especially influential during the first seconds of light onset. Peak PCA also shifts slightly towards lower wavelengths between five and fifteen seconds after onset in the case with darkness between light steps (dashed curve, shift from 525 nm to 500 nm, see also supplemental *Fig S10*), but not otherwise (maximum is stationary at 550 nm). PCA increases with *irradiance* (β_log10(irradiance)_ = +9.0% ±1.9 SE). When leaving *irradiance* out of the model, differences between the two settings of *Dark* stay mostly the same, whereas PCA sensitivity overall shifts slightly towards higher wavelengths (about 10 to 20 nm; supplemental *Fig S12*). Finally, settings with darkness between light steps allow for PCA analysis during the re-dilation phase, known as post-illumination pupil reflex (PIPR, *Fig 8 A2 Dark, and Fig 8 A2 “21^st^ second or 6^th^ second after lights-off”*). Interestingly, re-dilation is slowest for the shortest wavelengths, which is also the case when not controlling for *irradiance (Fig S12)*. Wavelengths above 600 nm were least influential on the PIPR *(Fig 8 A2 “21^st^ second or 6^th^ second after lights-off”*). The results for the protocols Long 1 and Long 2 of *Experiment III* are shown in *Fig 8 B* for better comparison with *Experiment II*, but are reported further below.

**Fig 8.**
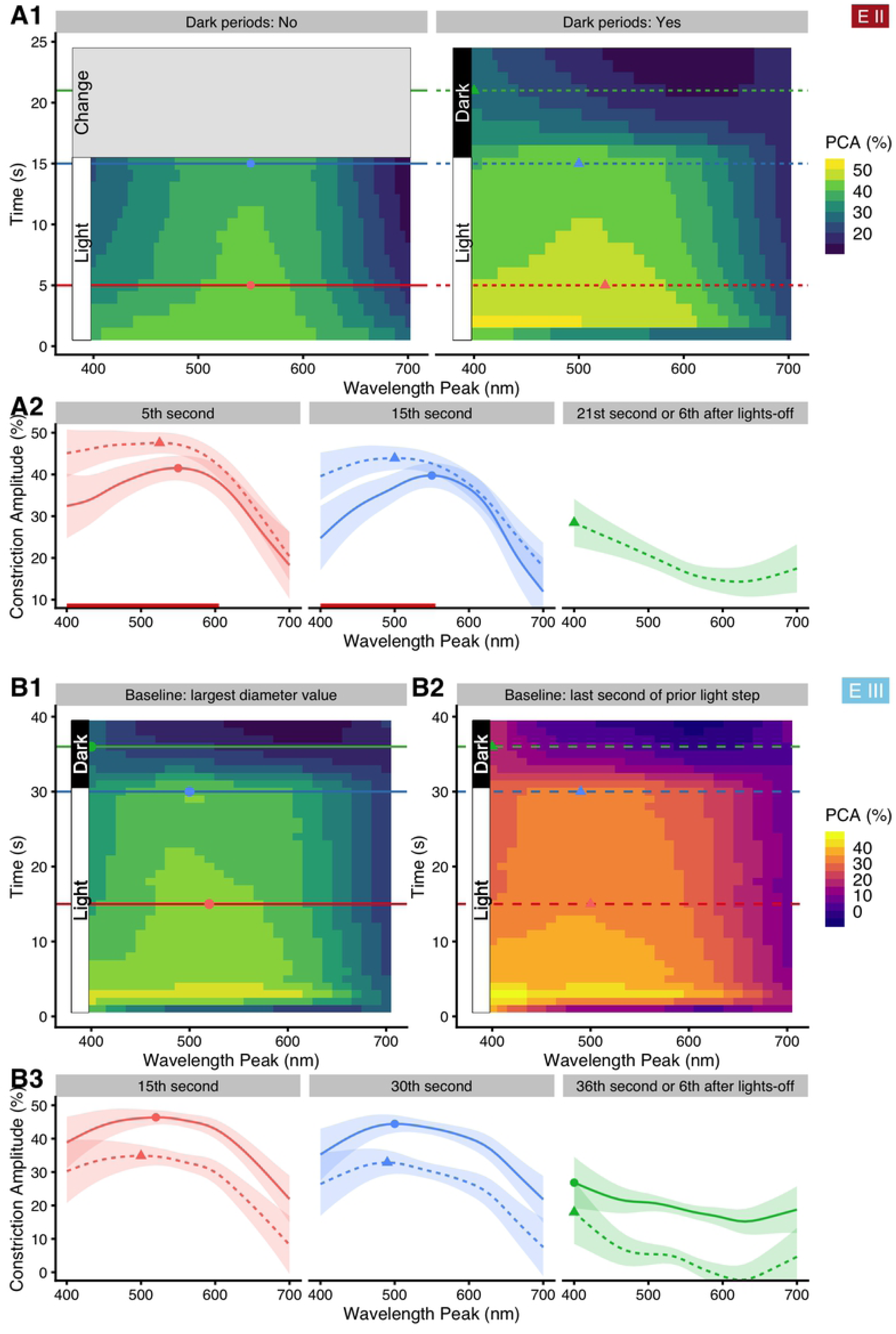
Interaction of *wavelength* with *time* in *Experiment II* and *III*. *Time* denotes the time (in seconds) since light onset, or light change to the respective wavelength. (A1) and (A2) show results from *Experiment II*, (B1), (B2), and (B3) those from *Experiment III* for the protocols *Long 1* and *Long 2*. (A1) False-color graph of model predictions for the PCA’s dependence on *wavelength* (x-axis) and *time* (y-axis) for settings with (right panel), or without (left panel), periods of darkness between changes of wavelength. All other predictors are held constant at their average value. Horizontal lines indicate where the respective traces shown in figure part (A2) are taken from; continuous curves refer to the absence of dark periods, dashed curves to their presence; filled circles and triangles mark the wavelength of the respective maximum PCA value of these traces. In the right panel of A1, PCA values after the 15^th^ second (i.e., in the Dark period) show a time-by-wavelength rendition of the post-illumination pupil reflex (PIPR). (A2) Model predictions for PCA vs. wavelength at three points in time. The right panel shows the wavelength dependency of the 6-second PIPR. Ribbons show the 95% confidence interval for the predicted means. Dotted lines with a triangle symbol represent the discontinuous setting with periods of darkness present between light steps; full lines with a filled circle show the continuous setting. The red horizontal line above the x-axis in the two left panels shows where the difference between the two settings is significant at the 0.05 level. (B1) Like (A1 right panel), but for *Experiment III*. Horizontal lines show where the respective traces shown in figure part (B3) are taken from, and points mark the wavelengths of the respective maximum PCA value of these traces. (B2) Like (B1), but with the PCA baseline taken from the last second of the respective previous light step. (B3) Like (A2), but dotted lines and triangles represent the scaling according to (B2), full lines and points according to (B1). The right panel shows the wavelength dependency of the 6-second PIPR.

#### Other dependencies

We tested PCA for an interaction of *wavelength* and *series*, which turned out not significant (p > 0.18). An influence of the prior wavelength on PCA was not significant either (p = 0.37). Furthermore, we used the average pupil diameter of the last second before the start of a light step as an alternative baseline value in *eq1*. We explored this variant as the response variable in the *wavelength-by-time* model for protocols using periods of darkness between light steps, where this PCA calculation is a valid alternative. As expected, *series* is no longer significant in this case (p = 0.31), since it is accounted for by the respective alternative baseline diameter. Irradiance is not significant either (p = 0.37). Otherwise, the model does not seem to behave differently in terms of *wavelength* or *time* (see supplemental *Fig S13*), other than that PCA values are, on the average, about 10% lower.

#### Illuminance

To analyze how well each type of illuminance can predict PCA in those protocols with periods of darkness between light steps, we constructed simplified standard linear mixed-effect models analogous to those used in *Experiment I*. The results were shown above in *Fig 7 B* (for comparison to *Experiment I*). As in *Experiment I*, they show that mesopic weighing predicts PCA best (R^2^_illu_= 0.55). Photopic functions do somewhat less well in comparison (R^2^_illu_ = 0.49 for V_10_ (λ), and R^2^_illu_ = 0.42 for V(λ)). Melanopic and rhodopic receptor functions work better than in *Experiment I* (R^2^_illu_ ≤ 0.31). Of the three cone functions, chloropic weighing led to the best results (R^2^_illu_ = 0.54), with the erythropic function second (R^2^_illu_ = 0.43). In contrast, cyanopic weighing was the only illuminance type that had no support by the model at all (p = 0.41, R^2^_illu_ < 0.01).

### Experiment III

#### Base model results

For the analysis of Experiment III, we analyzed PCA results for the protocols *Short 1* and *Short 2* in models separate from those for *Long 1* and *Long 2,* since the two protocol pairs differ conceptually. For those with one second of light followed by thirty seconds of darkness (*Short 1* and *Short 2*), PCA was calculated as the average PCA value of the sixth second after lights-off (or seventh second of the respective light step). The model shows neither support for an effect of *wavelength,* nor for one of *irradiance* (all p > 0.31).

For the two protocols with thirty seconds of light followed by nine seconds of darkness (*Long 1* and *Long 2*), PCA was calculated, as above, as the average PCA value of the last five seconds of light; the results were shown above in *Fig 5 C1*. *Wavelength* has support from the model (ΔAIC = 22). The predictor shows an inverted-U shaped curve, with a peak at 500 nm and only slightly lower amplitude at short wavelengths (*Fig 5 C1*). The effect of *series* is not significant (p = 0.54) (*Fig 5 C2*); PCA does not seem to depend on the stimulus’ position in the series when the stimulus duration is extended to thirty seconds. In contrast, PCA increases strongly with *irradiance* (β_log10(irradiance)_ = +19.7%, std. error 4.3). *Irradiance* has support (ΔAIC = 19) as part of the model (*Fig 5 C3*). Compared to other estimates of *irradiance*, the effect is rather large. See below for a discussion of a possible overcompensation. When removing *irradiance* as a predictor from the model (supplemental *Fig S15C*), the lower wavelengths become far less influential, with results similar to those in *Experiment I* (supplemental *Fig S15A*). 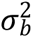 was 5.4% in *Experiment III*, which is the standard deviation of the random intercept by participant.

#### Time-course of the wavelength dependency

Analogous to the analysis in *Experiment II*, we used one-second averaged values in Experiment III to explore how PCA vs. *wavelength* changed over the time course of the light application. The results are shown below in *Fig 9* for the protocols *Short 1* and *Short 2*, and were shown above in *Fig 8B* for the protocols *Long 1* and *Long 2,* respectively. For the settings with one second of light, followed by thirty seconds of darkness (*Short 1* and *Short 2*), model diagnostics for PCA with the global pupil baseline per protocol were initially problematic, showing heteroscedasticity and skew in the residuals. The skew was eliminated by a logarithmic transformation of PCA and heteroscedasticity was reduced by a logarithmic transformation of *time*. *Time* is the time (in seconds) since light onset or light change to the respective wavelength. We also analyzed the data using the alternative PCA pupil baseline defined as the average diameter of the last second of the respective previous light step instead of the global baseline diameter taken across all light steps in one protocol. With the alternative baseline, model diagnostics were far more satisfactory compared to the global baseline, and only required logarithmic transformation of *time*, which is why this model is preferred. Both models used *adaptive splines* for accommodating the rapid changes during the first half of the light step compared to almost no changes during the second half. The results are shown in *Fig 9 A* and *Fig 9 B*, with exemplary sections in *Fig 9 C*. Both models predict that shorter wavelengths lead to a speed decrease in pupil re-dilation compared to longer wavelengths. The sixth-second of the PIPR (*Fig 9 C*, middle panel) shows a peak constriction at 490 nm for the preferred model, and 520 nm for the standard baseline. At the 15^th^ second (*Fig 9 C*, right panel), the peak has disappeared. The effect of *irradiance* is not significant in the preferred model (p = 0.88).

**Fig 9.**
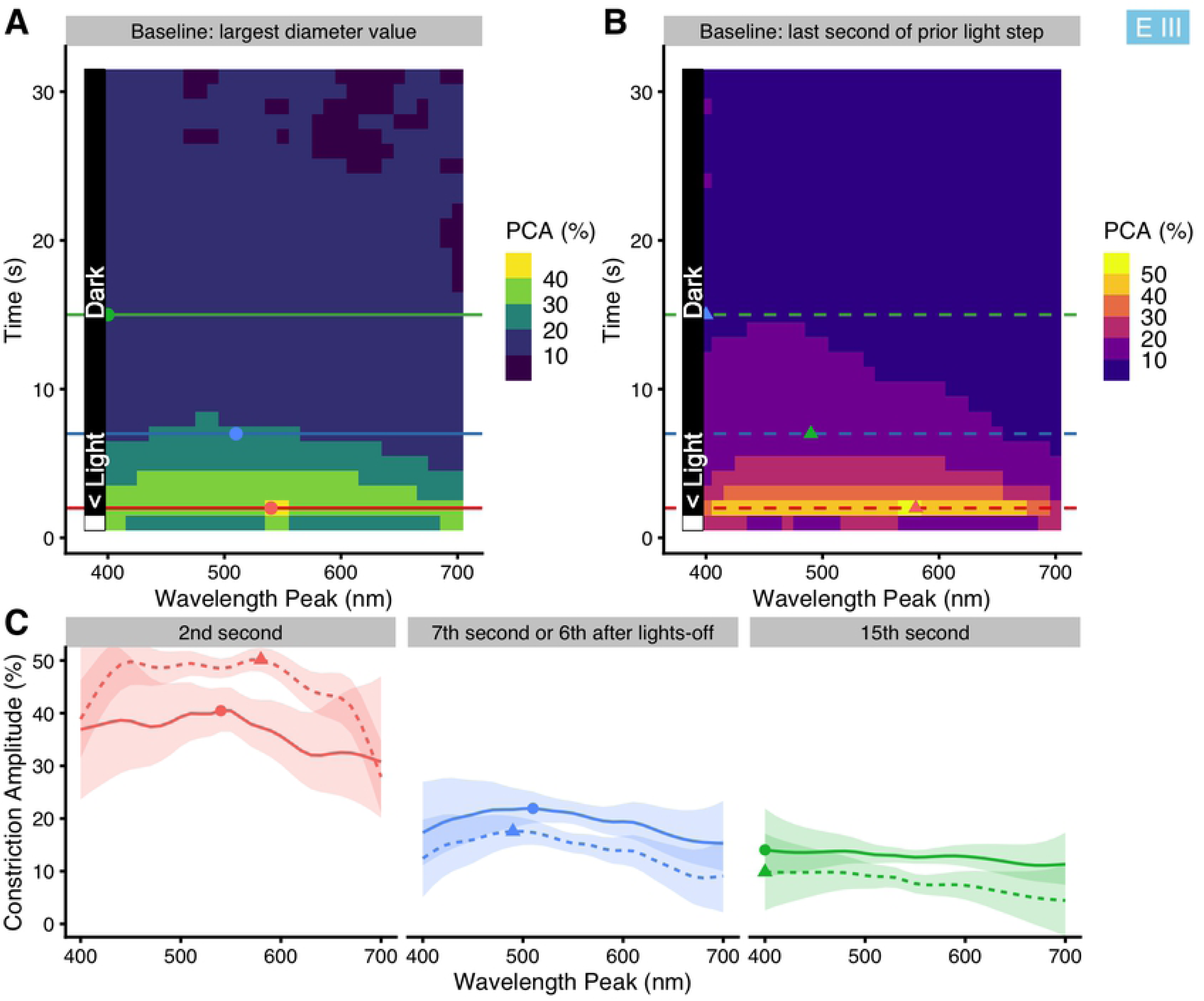
Interaction of *wavelength* with *time* in *Experiment III,* for protocols *Short 1* and *Short 2*. *Time* is the time (in seconds) since light onset or light change to the respective wavelength. (A) False-color graph of model predictions for the PCA’s dependence on *wavelength* (x-axis) and *time* (y-axis). All other predictors (basic model) are held constant at their average. Horizontal lines show where the respective traces shown in part (C) are taken from, and points mark the wavelengths of the respective maximum PCA value of these traces. (B) Like (A), but with a different measure of PCA where the PCA baseline is taken as the mean pupil diameter across the last second of the respective previous light step. Model diagnostics are superior to those for the Model in part (A). (C) Model predictions for the PCA vs. wavelength at three points in time after light-step onset: in the 2^nd^ second (red traces), the 7^th^ second (green traces), and the 15^th^ second (blue traces); the latter two cases are also the 6-second PIPR and 14-second PIPR, respectively. Ribbons show the 95% confidence interval for the predicted means. Dotted lines with a triangle represent the scaling according to (B), full lines with a filled circle according to (A).

For the settings with thirty seconds of light followed by nine seconds of darkness (*Long 1* and *Long 2*), model diagnostics for both baseline methods were satisfactory. The results for these models were shown above in *Fig 8 B1* and *Fig 8 B2*, respectively, with exemplary sections in *Fig 8 B3*. Both models predict that shorter wavelengths become more important with *time*, compared to longer wavelengths. With the standard baseline (*Fig 8 B1*), peak PCA is at 520 nm after 15 seconds, and at 500 nm at the end of the light application. With the alternative baseline (*Fig 8 B2*), peak PCA is at 500 nm after 15 seconds, and at 490 nm at the end of the light application. Re-dilation after lights-off is similar to that in *Experiment II* (*Fig 8 A*). PCA increases with *irradiance* (Baseline: largest diameter value; β_log10(irradiance)_ = +13.0% ±3.1% SE). When leaving *irradiance* out of the model, PCA sensitivity overall shifts slightly towards higher wavelengths (about 10 to 30 nm; supplemental *Fig S14C* and *S14D*), similar again to *Experiment II*.

#### Illuminance

To analyze how well each type of illuminance can predict PCA in the *Long* protocols (thirty seconds of light followed by nine seconds of darkness), we constructed simplified standard linear mixed-effect models analogous to those in *Experiment I* and *Experiment II*. The results were shown above in *Fig 7 C but are reported here*. As in the other experiments, mesopic weighing predicts PCA best (R^2^_illu_= 0.48). Photopic functions do less well in comparison (R^2^_illu_ = 0.45 for V_10_ (λ), and R^2^_illu_ = 0.40 for V(λ)). Curiously, melanopic and rhodopic receptor functions are less good predictors than in *Experiment I* or *Experiment II* (R^2^_illu_ ≤ 0.21). Of the three cone functions, chloropic weighing led to the best results (R^2^_illu_ = 0.44), with the erythropic function second (R^2^_illu_ = 0.42). In contrast, cyanopic weighing was the only illuminance type with no support by the model (p = 0.24, R^2^_illu_ < 0.01).

### Pooled data

To gain further insight into the interaction of *wavelength, chronotype,* and *time of day*, we repeated the analysis from *Experiment I* with pooled data from all experiments. Data from protocols *Short 1* and *Short 2* were excluded since these differ conceptually from the others. To partly account for differences between the experiments, we allowed *wavelength* to vary based on experiment type, which is a slight but acceptable oversimplification for *Experiment II* (see above). We further allowed *series* to change, based on the experiment. The interaction effect had strong support (ΔAIC = 163); results were shown above in *Fig 6 B*. Compared to the analysis based solely on the results of *Experiment I* (*Fig 6 A*), the basic patterns are similar for the evening and the afternoon. Before midday, however, *Larks* do not shift in wavelength sensitivity towards the shorter wavelengths as was the case in Experiment I. Further, the shift towards longer wavelengths for *Owls* is not as strong. Lastly, there is strong support of separate interaction effects (*wavelength, chronotype,* and *time of day)* based on whether or not there are periods of darkness between light steps (ΔAIC = 176), but the resulting model predictions do not lead to any insights beyond what is shown in *Fig 6*. Possible reasons for the differences between the models will be discussed in the next section. As a side note, the effect of *irradiance* was estimated to be an increase of PCA of 8.9% per order of magnitude (β_log10(irradiance)_ = +8.9% ±1.3 SE) irradiance increase.

## Discussion

### Base model results and time-course of the wavelength dependency

Our initial hypothesis was mostly based on findings from McDougal and Gamlin (11) and Gooley et al. (12). For a continuous series of monochromatic light steps, lasting 15 seconds each, we expected cones to contribute minimally after 10 seconds [11]. Furthermore, because of the whole series’ considerable length (just over 15 minutes), we expected sluggish ipRGC influences to manifest themselves [12]. Therefore, we believed wavelengths between 490 nm to 510 nm to have a considerable effect on pupillary constriction amplitude (PCA) ten to fifteen seconds into each light step. However, the results of *Experiment I* do not support this hypothesis (*Fig 5 A1*). The peak PCA around 540 nm is close to the peak of the long and middle-wave cone functions (L+M), which suggests strong cone influences. There is further support for an interaction of *wavelength* and *series* in *Experiment I* which indicates a slightly heightened PCA at, and shortly after, reaching the wavelength of peak ipRGC sensitivity, but the effect is small to begin with and disappears over the course of the protocol (supplemental *Fig S5*). Compared to other research [2-4, 8-12], only Alpern and Campbell (3) report similar results (532 nm peak for the 2.5 mm contraction threshold), while most other papers suggest a peak between 480 and 510 nm [2-4, 8-12], depending on experimental conditions.

There are certain differences and limitations when comparing our results with published literature. Firstly, as stated above, we used continuous light in the first protocols, i.e., participants were pre-adapted to light for all but the very first light step. In the literature, most stimuli are presented singly, with some period of dark adaptation in between [29]. This difference in adaptation also influenced the pupil baseline, as discussed above under *Materials and methods*. Secondly, due to our setup pupil reaction was evaluated at some fixed stimulus intensity, while, in contrast, in most publications stimulus intensity is varied to reach a fixed psychophysiological threshold (e.g., 50% constriction) [2]. Thirdly, we operated in the mesopic stimulus range (≤ 10 cd/m^2^), a limitation of our light source. Although this is well above the ipRGC-sensitivity threshold, stimuli of about one degree of magnitude higher intensity would have been desirable to elicit the strongest ipRGC reactions [17]. Lastly, differences in *irradiance* between wavelengths required that this variable is included as a predictor in the model, which is relevant in particular for wavelengths at and below 450 nm. Part of the intent of the second and third experiment was thus whether differences in our results from published literature are due to our setup, or due to our protocol, rather than data analysis. Since our initial hypothesis about ipRGC influence on the pupillary constriction amplitude (PCA) for a series of monochromatic light stimuli did not hold, we aimed to explore several dependencies with the available data and conducted two additional experiments aimed at specific questions arising from our experimental outcomes.

Light-step changes in the first experiment were small, with a wavelength shift of only 5 nm. We set up *Experiment II* to ensure that the wavelength dependency of the first experiment was not some artefact from the near-continuous sweep across the visible spectrum. Results revealed that this was not the case (compare the red trace in *Fig 5 B1* to the trace in *Fig 5 A1*). More importantly, results showed that short wavelengths had a far stronger influence when the light stimulus was discontinuous between light steps, compared to continuous light (*Fig 5 B1*, blue trace compared to the red trace). While the results for discontinuous light, i.e., with short periods of darkness between light steps, suggest a rod rather than an ipRGC influence, they agree far better with published findings [2-4, 8-12] than those from the first experiment. Regarding the time shift in wavelength dependency throughout each light step, the curve remains centered at around 550 nm with continuous light, but is shifted towards shorter wavelengths with discontinuous light (*Fig 8 A2*, full vs. dashed curves). In *Experiment III,* thirty seconds of light were followed by nine seconds of darkness. In this case, the last five seconds of light showed even more influence of short wavelengths than in the previous case (*Fig 5 C1*, compared to the cyan trace in *Fig 5 B1*), and the broad peak resembles the behavior shown by Mure et al. (10) for the *0–30 second* condition. The shift in wavelength dependency is also more pronounced in *Experiment III* than in *Experiment II* (*Fig 8 B1 and 8 B3*, compared to *Fig 8 A*). In summary for our results, the cone influence fades in favor of rod influence for discontinuous series of long-lasting monochromatic light steps, agreeing with findings from McDougal and Gamlin (11). However, cone influence does not fade for pre-adapted participants as part of a continuous series of light steps, which to our knowledge has not been reported before. Mure et al. (10) showed changes in wavelength dependency occurring with pre-exposure, but this applied for high-intensity stimuli with several minutes of darkness in between. Joyce et al. (55) showed the effect of short-term light adaptation on the PIPR, but not on the light-adapted pupil itself. The continued bleaching of the sensitive rod photoreceptors might desensitize the rod channel, leaving only the changes in cone input as contributors to PCA. While this mechanism cannot be deduced from our data, it seems a plausible explanation and would result in the shown spectral dependency. It was suggested by McDougal and Gamlin (11), however, that the rod channel remains relevant over minutes of continuous light. Instead, we saw that the spectral dependency of the pupillary reaction to monochromatic light is similar to the spectral dependency of polychromatic light [14] when monochromatic stimuli are applied under pre-adaption. In vision, interaction effects of rod and cone inputs that depend on the adaptation state of receptors are well known (for a review, see Zele and Cao (56)). In any case, and despite the mesopic experimental conditions, we would have expected some form of ipRGC contribution, which was not apparent, however. Spitschan et al. (57) showed an S-cone opponency to ipRGC input in the pupillary light response, where increasing S-cone stimulation reduced PCA, all other receptor types being stimulated at a constant rate. If cones remain more relevant under pre-adaptation to light compared to prior darkness, as our results suggest, S-cone opponency might help to explain the reduced short-wavelength sensitivity in the presence of relevant ipRGC input to pupil control. IpRGC influence might still be visible in our data, however, as discussed below in *Chronotype and time of day*.

By their presence of a dark period after light stimulation, *Experiment II* and *III* allowed for the analysis of the post-illumination pupil response (PIPR). Surprisingly, the shortest wavelengths led to the PIPR’s slowest pupil re-dilation with 15 seconds of prior light in *Experiment II* (*Fig 8 A,* supplemental *Fig S12*), and even still with thirty seconds of prior light in *Experiment III* (*Fig 8 B,* supplemental *Fig S14*). Gamlin et al. (8), in contrast, reported that a Vitamin-A_1_ pigment nomogram with peak sensitivity at 482 nm fitted their data closely for the PIPR of two human subjects after only ten seconds of light. However, when looking at the individual data points for *log relative sensitivity* of the human pupil from that study below 482 nm, i.e., at 452 and 473 nm, their sensitivity does not seem to drop far below that of their closest neighboring data point (493 nm). Park et al. (17) showed that the ipRGC influence is best seen during the sixth second of re-dilation after a one-second light stimulus (compared to a ten-second stimulus in that study). The protocols *Short 1* and *Short 2* in *Experiment III* were constructed accordingly (results shown in *Fig 9* and supplemental *Fig S14*). We see a peak PCA at around 490 nm, which would fit the ipRGC sensitivity maximum for a young adult [28]. These results agree with a publication by Adhikari et al. (58). That study showed an apparent peak sensitivity between 464 and 508 nm (they fitted a Vitamin-A_1_ pigment nomogram with peak sensitivity at 482 nm), with an order of magnitude lower relative sensitivity at the 409-nm wavelength stimulus. After the sixth second in our results, however, sensitivity further shifts towards shorter wavelengths. Thus, our results on the PIPR do not contradict published literature but rather suggest a stronger influence of shorter wavelengths than previously thought, below the ipRGC peak at around 490 nm. A possible mechanism might be an S-cone influence on the pupil [59, 60], which has been shown to influence other ipRGC-dependent effects, such as circadian alignment in mice [61] or, very recently, acute Melatonin-suppression in humans [62]. Circumstances for these additional influences include low light levels and comparatively short stimulus times (less than half an hour), which would fit our setup. The S-cone influence on melanopic effects is not undisputed, however, as Spitschan et al. (63) found no evidence for the influence in acute neuroendocrine and alerting responses. A more far-fetched, but possible, mechanism is the influence of other opsin types in the retina, such as neuropsin (Opn5), which have been found in the retina of vertebrates (for a review see Guido et al. (64).

Regarding the series effect, there seems to be a cumulative effect in *Experiment I* and *II* for the first 18 steps (*Fig 5 A2 and 5 B2*), where PCA at first increases with each consecutive light step. The downward slope in *Experiment I* then after about five minutes (*Fig 5 A2*) might be the result of desensitization, where the small differences in wavelength between light steps are not pronounced enough to counter pupillary escape over time. In *Experiment II,* wavelength differences between light steps were larger, and PCA after step 18 did not noticeably increase further, nor decrease as in *Experiment I* (*Fig 5 B2*). Data analysis for *Experiment I* had suggested an interaction of *series* with *wavelength*. However, further analysis had shown that this effect is of little relevance to the interpretation of the data (supplemental *Fig S5*) and does not suggest ipRGC influence as initially assumed in *Materials and methods*. We also found no indication of influences from the prior wavelength to the current wavelength in *Experiment II*.

The estimated effect of *irradiance* varied between experiments and depended on whether five-second (base model) or one-second averages (time-shift model) of PCA were analyzed. While this is plausible insofar as the experiments differ conceptually, it is still worth discussing the estimates, summarized in *Table 1*. Except for the *Short* protocols in *Experiment III,* i*rradiance* was a significant predictor in all cases. In the *Short* protocols, the *irradiance* differences between the light steps were seemingly not big enough to influence re-dilation significantly. For the other cases, the estimated value is smallest for *Experiment I,* which is plausible since the changes in *irradiance* between consecutive light steps are the smallest across the experiments. In *Experiment II*, the effect of *irradiance* is similar between the base model and the time-shift model. In *Experiment III*, the estimated effect of *irradiance* for the base model is rather large compared to the other experiments and the time-shift model. The estimate for the time-shift model in *Experiment III* is closer to estimates in *Experiment II*. We believe the estimate for the base model of *Experiment III* to be unlikely high and see no theoretical basis for the large difference to the other estimates. This would mean that the *wavelength* effect of the base model in *Experiment III* overcompensates for *irradiance*. Therefore, the time-shift model is to be preferred to the base model, but the interpretation of *wavelength* as discussed above does not change in a relevant manner (compare *Fig 8 B3: 30^th^ second* to *Fig 5 C1*). For the model from pooled data of all experiments, estimates for *irradiance* were close to *Experiment II.* Setting the base model in Experiment III aside in favor of the time-shift model (see above), the differences between and within experiments stay broadly the same regardless of the estimated value of *irradiance*, thereby retaining the interpretations above regarding PCA’s dependency on *wavelength*.

**Table 1.** Model estimates for irradiance across all experiments.

|  | Base model | Time-shift model |
| --- | --- | --- |
| Experiment I | 6.5±1.4% | - |
| Experiment II | 10.7±3.4% | 9.0±3.4% |
| Experiment III | 19.7±4.3% | 13.0±3.1% |
| Pooled Data | 8.9±1.3% | - |

Values denote model estimates of $\beta_{\log_{10}(\text{irradiance})} \pm \text{SE}$ , which indicates the effect of $\log_{10}$ transformed irradiance on the pupillary constriction amplitude (PCA). The base model uses the average PCA of the last five seconds of light during each light step, the time-shift model uses one-second means of PCA along the time course of each light step

Finally, regarding the base models, the PCA-by-participant random effect in all three experiments show the large interindividual differences in pupil reaction, even after normalization. These interindividual differences are often remarked in other publications, as summarized by Kelbsch et al. (29). Depending on the experiment, the average PCA is predicted by the model to vary by about 20% between two extreme participant cases (0.025 to 0.975 percentile difference).

#### Chronotype and time of day

The models from *Experiment I* and pooled data across all experiments strongly suggest the presence of an interaction effect that influences PCA’s *wavelength* dependency by an individual’s *chronotype* and the measurement’s *time of day*. Specifically, early chronotypes, or *Larks*, before noon seem to have a heightened sensitivity to shorter wavelengths (*Fig 6 A1* and *B1*). In contrast, late chronotypes, or *Owls*, before noon seem to have a slightly lowered sensitivity to shorter wavelengths. These chronotype-specific differences in sensitivity before noon are called into question by the model constructed from pooled data across the experiments (*Fig 6 B1* and *B2*). Both models agree, however, that in the early evening (6:30), the wavelength of maximum sensitivity shifts to longer wavelengths for the *Larks*. For them, across the day the maximum sensitivity shifts from the longer to the shorter wavelengths, opposite to the behavior in *Larks*. In general, the shifts are predicted to be stronger for the more pronounced or extreme chronotypes, compared to moderate types. For *Neutral* types, the shift is predicted to be minimal. We also see the chronotype bias in *Experiment I* in the daytime scatterplot (*Fig 4 B,* the green dots show *Experiment I*), mostly for *Owls* during the first half of the day, which is very likely due to *Owls*’ later preferences. We pooled the data across all experiments in the hope to gain more insight from the available data. The pooling adds fifty protocol-runs to the eighty runs of *Experiment I* and leads to a more well-rounded distribution of chronotypes across the day (*Fig 4 B,* spread of the dots of all three colors). The pooling also introduces two new dependencies that might influence the interaction and need to be considered. Firstly, *Experiments II* and *III* were conducted decidedly later in the year, compared to *Experiment I* (*Fig 4 C*). The thereby introduced general time-shift by the shortened photoperiod should not be too influential according to a recent paper by McHill et al. (65). In that study, the time shift in dim-light melatonin onset was less than half an hour across three months from February to May. However, the shortened photoperiod might lead to changes in the possible mechanism underlying the interaction effect ([66], see below). Secondly, whether the light stimulus in the experiments is continuous or discontinuous makes a difference in wavelength dependency (*Fig 5 B1)*, and likely also with respect to the involved receptor types (see above for the discussion of the base model). Different receptor inputs likely lead to differences in the interaction, which is supported by a model differentiating the interaction of *wavelength, chronotype,* and *time of day* based on whether or not there are periods of darkness between light steps. We slightly prefer the simpler model constructed from *Experiment I*, because while pronounced chronotypes before noon are lacking (Fig 4 B, green points), it avoids the likely confounding influences from the pooled data mentioned above. It ultimately makes little difference which of the models is preferred, as both indicate the presence of a similar interaction of *chronotype* with *time of day* on the wavelength dependency. The precise nature of this dependency is beyond this study’s scope. It will have to be approached in a controlled experiment, with measurements for all chronotypes evenly distributed across all times of day.

Circadian effects on the pupil response were reported before and we will discuss two relevant publications in the context of our results. While the focus in those studies is different from ours, we can still compare how their results on circadian ipRGC input line up with our circadian effect. This is under the assumption, that our circadian effect is moderated by ipRGC input, as discussed below. It is further important that, in those studies, the effects of chronotype are adjusted for by looking at participants’ internal time or circadian phase, instead of at external time. While this avoids confounding from differences in internal time between participants, it also makes it hard to determine how much of a difference the chronotype signifies. The MEQ, which we used to assess chronotype, does not describe the participant’s phase of entrainment of, but rather their time of preference [67]. And while there are correlations between the phase of entrainment and the MEQ score, an exact phase shift cannot be deduced from the score. Therefore, assumed time-shifts between chronotypes in the following section are to be taken tentatively.

Zele et al. (23) showed a circadian response of ipRGC input to the PIPR on eleven participants in a 20–24h laboratory experiment, independent of external light cues. They also investigated circadian changes in the pupil reflex during their ten second light stimulus, but found only growing effects from fatigue owing to the long experiment. IpRGC input was evaluated in that study through the reduction of normalized pupil diameter after lights off (PIPR), compared between a blue and a red monochromatic light stimulus (peak wavelength at 488 nm and 610 nm, respectively). The normalized pupil diameter started to decrease about five hours before the evening melatonin onset (DLMO) for the red stimulus, and three hours prior to the DLMO for the blue stimulus, thereby implying a shift in wavelength dependency. The normalized pupil diameter reached its minimum shortly after the DLMO for both wavelengths; the overall period with a reduced pupil diameter lasted longer for the red than the blue stimulus. As mentioned above, any effect on the circadian ipRGC input due to chronotype is not discernible from these results, since the study centered participants’ rhythms on their respective points of melatonin onset, thereby eliminating chronotype differences. However, we would expect the effect of chronotype on melatonin onset to be a pure time-shift of a few hours [68]. Translating these findings to our context, we would expect a fixed relationship between PCA curves of comparable wavelengths across the chronotypes, with a general time-shift by about 1.5 to 2 hours between *Owls* and *Neutral* types, and another shift between *Neutral* types and *Larks*. As shown in *Fig 10,* our analysis rather suggests a crossed response of PCA depending on wavelength and chronotype (cut-off at 8pm, after which we did not collect data). We thus do not believe that the circadian effect described by Zele et al. (23) is the same as that in our data. Since that study focused on the afternoon and evening, the comparison to our data does not change depending on whether the data from *Experiment I* or the pooled data are used.

**Fig 10.**
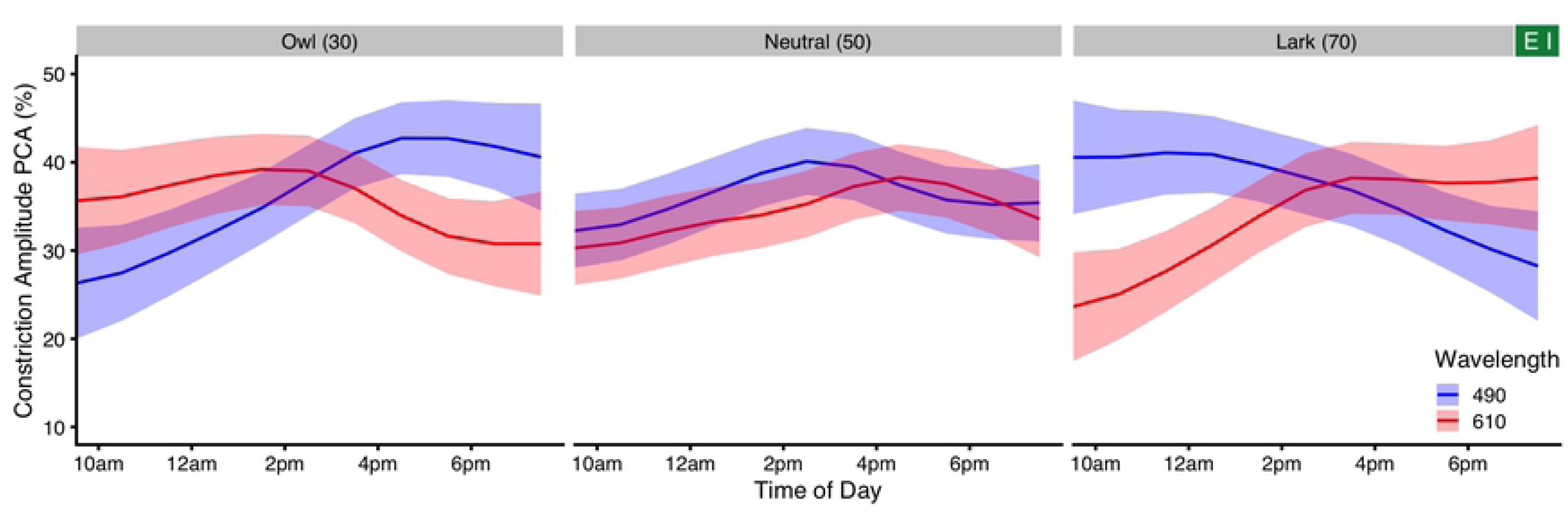
Select changes of PCA across the day in *Experiment I*. Model predictions for the PCA vs. time of day for three chronotypes (left to right), and two wavelengths (blue trace 490 nm and red trace 610 nm), that were chosen for comparison with the studies by Zele et al. (23) and Munch et al. (22). Ribbons show the 95% confidence interval for the predicted means.

Another study, by Munch et al. (22), also reported circadian effects on the PIPR. For ten subjects throughout two 12-hour sessions, the 6-second PIPR was assessed after 1-second and after 30-second light stimuli. The authors adjusted the circadian phase in a similar manner to Zele et al. (23). The normalized 6-second post-stimulus pupil size showed a circadian behavior in the short-wavelength, but not the long-wavelength, stimulus. Translating circadian phase and normalized pupil size to our context, PCA for blue stimuli would be highest before noon und decrease afterwards, with *Owls* having a later shift compared to *Larks*. Again, these predictions do not fit our data.

In both of the abovementioned studies, we expect the effect of chronotype to be an effect of time-shift of a few hours [68]. However, our model suggests something different, namely an interaction similar to the so-called synchrony effects of chronotype reported in the context of cognitive performance in education [69, 70], or of physical performance in athletes [71]. Goldstein et al. (70) define chronotype synchrony as the state in which the time of optimal performance is equal to the time of preference of the chronotype, e.g. morning for *Larks* and evening for *Owls*. Our model from *Experiment I* predicts such a heightened sensitivity for short wavelengths at the time of preference for the chronotypes. This connection might even go one step further. It is known that the time-of-day is highly relevant for the magnitude and sign of a circadian phase shift for a given stimulus [72] (for a review of known influences, see Prayag et al. (73)). Furthermore, Roenneberg and Merrow (74) argue that chronotype is not only the result of non-24-hour internal periods but also of the individual’s photosensitivity. Following this reasoning, heightened sensitivity to shorter wavelengths at circadian intervals could be part of a reciprocal system to strengthen or even cause a chronotype, rather than being simply a correlation. *Reciprocal* meaning that some mechanism – like an internal period of other than 24h duration – might predetermine a circadian type. Heightened sensitivity to wavelengths for a circadian shift at times of preference (e.g., towards the evening in *Owls*) means one would need more light outside of the preferred time to shift towards a different chronotype. Conversely it would need less light for strengthening the rhythm at the time of preference, thereby solidifying the chronotype. While causation cannot be determined from our study, intraocular mechanisms have been described to allow for such an effect. In general, rod and cone sensitivity follow a circadian rhythm [75, 76], with melatonin and dopamine as key actuators. Furthermore, ipRGCs are intricately and reciprocally connected to dopaminergic amacrine cells (for a review in the context of myopia, where dopamine plays an essential role, see Stone et al. (77)). Through this connection, ipRGCs can attenuate the outer retinal light adaptation in mice [78]. This system is involved in seasonal and circadian regulation of photosensitivity [66]. Other seasonal changes, like that of human color perception [79], might also be connected to this mechanism. Since ipRGC sensitivity is the basis of the mechanism, the shift towards, or away from, the short wavelength spectrum is plausible. Whether the mechanism operates in a chronotype-dependent manner, as shown above, is open for evaluation.

#### Sex and age

We saw strong support for the model which included *sex* as a predictor. There were no differences in dependence on *wavelength* related to *sex*. However, women were estimated by the model to have a lower average PCA. Chellappa et al. (21), in contrast, found sex differences in light sensitivity. In their study, men showed a stronger response to blue-enriched light in the sleep EEG (NREM sleep slow-wave activity) and for vigilant attention, and had higher brightness perception during blue-enriched light. Men also found blue-enriched light preferable to non-blue-enriched light, contrary to women. The stronger average PCA for men in our study would align with a higher brightness perception for men per se, but unlike in Chellapppa et al.’s study was not limited to short wavelengths. Since men – in general – are more likely to be late chronotypes compared to woman [80], and Chellappa et al. (21) did not mention whether they stratified their sample for chronotype, their reported effect might also have been caused, in part, by a chronotype effect. That study was performed in the late evening, where our interaction effect of chronotype with time of day and wavelength suggests a heightened sensitivity for short wavelengths (see above).

*Age* was estimated to influence PCA, but there was no indication of an interaction with *wavelength*. The latter agrees with a publication by Rukmini et al. (81), where pupillary responses to short-wavelength compared to long-wavelength light were independent of age (in a comparison of two age groups: 21–30, and ≥50). Depending on whether or not chronotype and time of day were included in our model, PCA was predicted to rise steadily with age (*Fig 5 D1,* supplemental *Fig S6 A*), or reach a plateau at about age 25 (supplemental *Fig S6 B*). According to Rukmini et al. (81), pupillary responses were reduced in older participants. *Age* was also part of the unified model from Watson and Yellott (1). However, the latter focused on pupil diameter itself, rather than the (relative) change in diameter, i.e., PCA. We can still infer from their predictions that PCA would likely shrink with age, because there is a natural lower limit to the pupil diameter, thereby decreasing PCA with an ever-shrinking upper limit of pupil diameter. Conversely, Daneault et al. (32) found no significant age-related differences in pupil constriction, even though pupil size itself decreased with age. Adhikari et al. (82) showed that the relative peak constriction did not change with age, that measure being closely related to our dependent variable PCA. While not significantly correlated with *age*, the trends in their study for both blue and red stimuli show a slight positive slope. In summary, literature reports show a heterogeneous picture of the effect of age on PCA. Our age range was small by design and focused on participants below the age of forty. Accordingly, the effects of *age* in our study were small and appear compatible with any of the studies mentioned above, thereby contributing little for this specific aspect.

#### Illuminance

At present, there are nine distinct measures of illuminance in common use to describe visual and alpha-opic (i.e., single receptor-dependent) effects of light. Scotopic illuminance and rhodopic equivalent daylight illuminance count as two, but both use the same rhodopic action spectrum. Many publications use V(λ)-derived luminance [1] (in cd/m²), but since the spatial stimulus characteristics are unchanged across our experiments (Ganzfeld conditions), luminance and illuminance are interchangeable in terms of comparisons between the types of illuminance. Adrian (5) argued that mesopic weighing was sufficient to explain apparent wavelength-dependency shifts towards the shorter wavelengths (under mesopic conditions). Accordingly, Adrian explained the mechanism behind this shift as purely S-cone and rod-based. We now know that Adrian (5) was incorrect in his assumptions about the sole mechanism and that there is additional input from ipRGCs in the pupillary system, e.g. from the work of Gamlin et al. (8) or Zaidi et al. (9). However, the appeal of the mesopic weighting compared to others is easy to see across all three of our experiments (*Fig 7*), where the mesopic illuminance best predicted PCA, thereby considering all cone and rod input. The single-opsin weighting functions for rhodopsin or melanopsin perform far less well in all cases.

In summary, our findings are relevant in the context of arguments made by Spitschan (13) and Zandi et al. (14) about the limited applicability of luminance-based pupil models [1] in the context of ipRGC inputs. At least for our experimental setup of monochromatic and mesopic light stimuli of changing wavelength, mesopic weighting was best at predicting pupil constriction. Conventional photopic (il)luminance-based approaches performed still well and, compared to single-opsin based functions, were preferable to predict pupil constriction. Finally, Spitschan et al. (57) reported an S-cone opponency, where the pupil would curiously increase in diameter with increasing stimulation of S-cones. Since our setup did not account for the constant stimulation of all other receptor types as that study did, any opponency effect is expected to be decreased in our study. The cyanopically weighted equivalent of daylight illuminance (S-cone weighing) was not significantly correlated with PCA in any experiment here. The cause is not known, but the results are compatible to a zero-sum of opposing inputs.

## Conclusion

The spectral dependency of pupillary reactions to monochromatic light is well understood for isolated stimuli, but less so for scenarios with preadaptation to light. We looked at the pupillary constriction amplitude (PCA) in protocols of monochromatic light of periodically changing wavelength and found very little influence of short wavelengths. A second experiment showed that this effect originated in the continuous application of light and that the effect was present over the whole time-course of each wavelength. Our results for continuous monochromatic light and literature findings from polychromatic light share a similar wavelength dependency, compared to singularly presented monochromatic light. Furthermore, the 6-second post-illumination pupil response (PIPR) to isolated light stimuli, for two out of three durations showed that the slowest re-dilation happens for the shortest wavelengths, well below peak ipRGC sensitivity, implying the presence of different, or at least additional, receptoral influences to the PIPR. Pupillary reactions to isolated light stimuli and the PIPR to short light stimuli behaved very much as would be expected from published literature. This strengthens our belief that our results are valid despite some technical limitations as described above. We further showed that mesopic illuminance was the best measure of illuminance to explain the pupil reaction. We found dependencies of *sex* and *age* on the pupillary response through the exploration of several covariates, but mainly an interaction of *chronotype* and *time of day* with *wavelength*. The interaction effect implies a modulation of wavelength-sensitivity with a heightened sensitivity to shorter wavelengths at the time of chronotype preference. This circadian effect further seems to manifest itself differently, depending on whether light is continuous or discontinuous. The effect could be linked to a mechanism that strengthens an individual’s chronotype at the current time of preference. If the effects are reproducible in a controlled experiment, they might even as a form of marker for the individual’s chronotype.

## Acknowledgements

We thank Jorge Alberto Gutiérrez Alvarez, Theresa Scherzer, Daniel Philipp Setzensack, and Regina Heiß for their help with data collection as part of their respective thesis work. We also thank Moritz Faust for his support on data collection and questionnaire digitalisation.

## Supporting information

**S1 Fig. PCA depending on *wavelength* for different dark adaptation periods (DAPs).** Traces show the average, LOESS-smoothed PCA vs. wavelength for two subjects, each repeating all shown protocols three times. DAP varied between 1.5, 3, and 15 minutes (dark to medium grey traces). For comparison, results of a second, consecutive run (also performed three times) of the first protocol is shown, as is the *Down* protocol (light grey). Points show the PCA values from which the traces were constructed. More information is found in *Materials and methods*.

**S2 Zip File. R-Markdown script results for calculating PCA from measurement data.** Zip file containing nine html files with results from R-Markdown scripts. The scripts are one each for the nine protocols used across the experiments. Since the raw measurement data are not part of the provided supplements, we included one exemplary result file for each protocol instead of the script itself. These html files show the code, as well as the output for the example participant.

**S3 Microsoft Excel File. Irradiance measurements and illuminance values for the range of light stimuli.** Microsoft Excel file containing two worksheets. The worksheet ‘Measurements’ contains the irradiance measurements in a resolution of 1 nm. The worksheet ‘Illuminance’ contains all types of illuminance values described in the main text and irradiance. All displayed values are based on spectral irradiance measurements with a field-of-view restriction according to the CIE S 026 standard [28]. In our case, these measurements are 24% lower than those of the unobstructed sensor diffusor.

**S4 Zip File. R-Markdown script used for statistical analysis and graphics generation.** Zip file containing several files necessary to replicate the statistical analysis and generation of graphics. Besides some external tables, an R-function file, and three pictures to mark the respective experiment in graphs, the zip file consists of seven R-Markdown scripts. One of these is used to set up the data prior to analysis. The necessary data can be downloaded from the *Open Science Framework* [35]. Five of the R-Markdown files are for analysis of *Experiment I, II, III Short, III Long*, and the pooled data. The last file is for graphics generation.

**S5 Fig. PCA depending on wavelength with and without interaction in *Experiment I*.** Model prediction for models with (dashed lines), and without an interaction effect (solid lines) of *wavelength* and *series,* across the series. The number above each plot indicates the series number. Colored dots represent raw PCA values at the respective series number and wavelength, their color indicates the respective protocol. The positions of points on the x-axis indicate where changes between the two models (lines) should be evaluated. For the model with the interaction effect, it seems as though PCA is increased after reaching the ipRGC peak sensitivity (at about 490 nm for a young adult), but not strongly and not for long.

**S6 Fig Influence of *age* in *Experiment I*.** (A) Model predictions how PCA depends on *age*, when all other predictors (basic model) are held constant at an average level. (B) Like (A), but for the model with all other dependencies in *Experiment I* included. Here, *age* seems to affect PCA with a more continuous rise over the age range.

**S7 Fig. GIF Animation of PCA depending on *wavelength*, *time of day*, and *chronotype* in *Experiment I*.** Model predictions show time-of-day values at half-hour points from 8:30 am to 7:30 pm. Left: False-color model predictions for hourly PCA values depending on *wavelength* (x-axis) and *chronotype* (y-axis), when all other predictors (basic model) are held constant at an average level. Horizontal lines show where the traces from the right image are taken from. Right: Model predictions for hourly PCA values depending on wavelength for three chronotypes: *Larks* (green traces, CT score 70), *Owls* (red traces, CT score 30), and *Neutral* types (blue traces, CT score 50). Ribbons show the 95% confidence interval for the predicted mean values.

**S8 Fig. PCA depending on *wavelength*, *time of day*, and *chronotype* in *Experiment I*.** False-color contour-line graphs of PCA model predictions for several wavelengths between 430 and 670 nm (color scale shown in the inset at the upper right corner of each plot). All plots are scaled equally. Each plot visualizes PCA depending on *time of day* (clock time in 24h values, x-axis) and *chronotype* (higher values are morning types, lower values evening types). Plots were created with the *vis.gam()* function in R, with the *too.far* argument set to 0.1. The *too.far* argument excludes grid points from the plot, when points are not represented by variable combinations close enough to actual data. Thereby, *too.far* is a measure of accepted extrapolation, scaled from 0 to 1.

**S9 Fig. Basic model predictions for unscaled changes in pupil diameter, and raw pupil diameter, in *Experiment I*.** Model predictions for response variables other than PCA, versus three main predictors (*wavelength, series,* and *irradiance*), when all other predictors are held constant at their average level. Traces show the model prediction for the mean, ribbons its 95% confidence interval. Red dotted lines show particular predictor values for the plotted relationship. (A / D) Response vs. stimulus peak wavelength. (B / E) Response vs. series number of light steps. (C) Response vs. stimulus irradiance. The x-axis scaling reflects the logarithmic transformation of irradiance.

**S10 Fig. GIF Animation of pupillary constriction amplitude (PCA) depending on *wavelength* and *time* in *Experiment II*.** Left: False-color model predictions for PCA depending on *wavelength* (x-axis) and *time* (y-axis), for settings with, or without, periods of darkness between wavelength steps. All other predictors are held constant at an average level. Horizontal lines show where the traces from the right image are taken from. Right: Model predictions for the PCA vs. *wavelength* over *time*. Ribbons show the 95% confidence interval for the predicted means. Blue lines represent the setting when periods of darkness are present between light steps, red lines when not. The red horizontal line above the x-axis shows where the difference between the two settings is significant (5% level).

**S11 Zip File. Html file with 3-D visualizations of pupillary constriction amplitude (PCA) depending on *wavelength* and *time* in *Experiment II*.** The zip file contains three files. One html file with a scalable and rotatable diagram of PCA vs. wavelength and *time*. An html viewer, or browser with support for html widgets, is needed. The other two files are snapshots from the html file for each condition of *Dark*. Color mapping is according to the z-axis (PCA) for better visibility.

**S12 Fig. Interaction of *wavelength* with *time* in *Experiment II*.** (A) False-color graph of model predictions for the PCA’s dependence on *wavelength* (x-axis) and *time* (y-axis) for settings with, or without, periods of darkness between wavelength steps. All other predictors (basic model) are held constant at their average. Red dots show the peak PCA value for each second, the value right next to it the respective peak wavelength. Green dots and values show the respective trough for PCA. The trough is at 700 nm where green dots are not shown. (B) Like (A), but for the model without *irradiance* as predictor.

**S13 Fig. Interaction of *wavelength* with *time* in *Experiment II,* for protocols with darkness between light steps.** (A) False-color graph of model predictions for the PCA’s dependence on *wavelength* (x-axis) and *time* (y-axis). All other predictors (basic model) are held constant at their average. Horizontal lines show where the respective traces shown in part (C) are taken from. (B) Same as in (A) except that the baseline for PCA calculation (*eq1*) is taken from the last second of the respective previous light step. (B) The graph serves to check whether and how the single baseline value per protocol (shown in A) changes the model prediction. (C) Model predictions for the PCA vs. wavelength at three points in time after light step onset: in the 5^th^ second (red), 15^th^ second (blue), and 21^st^ second (green); the latter case is also the sixth seconds after lights-off. Ribbons show the 95% confidence interval for the predicted means. Full lines represent PCA values when baseline pupil values are taken from each protocol’s largest pupil diameter, dotted lines when baseline pupil values are taken from the last second of the respective prior wavelength-step.

**S14 Fig. Interaction of *wavelength* with *time* in *Experiment III*, for models with, or without, irradiance as predictor.** (A) False-color graph of model predictions for the PCA’s dependence on *wavelength* (x-axis) and *time* (y-axis) for the protocols with one second of light, followed by thirty seconds of darkness, and with *irradiance* as the predictor. All other predictors (basic model) are held constant at their average. Red dots show the peak PCA value for each second, the value right next to it the respective peak wavelength. Green dots and values show the respective trough for PCA. The trough is at 700 nm where green dots are not shown (e.g. between 2 and 30 seconds). The baseline for PCA calculation (*eq1*) is taken from the last second of the respective previous light step, since this baseline led to a preferable model in terms of model diagnostics. (B) Like (A), but for the model without *irradiance* as the predictor. (C) Like (A), but for protocols with thirty seconds of light, followed by nine seconds of darkness with irradiance as a predictor. The baseline for PCA is as described in *Materials and methods*. (D) Like (C), but for the model without *irradiance* as the predictor.

**S15 Fig. Model predictions for models without *irradiance* as a predictor.** Model predictions for pupillary constriction amplitude (PCA) as depending on *wavelength*, when all other predictors are held at an average, constant level. Traces show the model prediction for the mean, ribbons its 95% confidence interval. Dotted lines show the respective peak. (A) through (C) show dependencies in *Experiment I*, *II*, and *III*, respectively. They can be compared to the model results which include irradiance in *Fig 5 A1*, *B1*, and *C1*.

